# A First-principles Approach to Large-scale Nuclear Architecture

**DOI:** 10.1101/315812

**Authors:** Ankit Agrawal, Nirmalendu Ganai, Surajit Sengupta, Gautam I. Menon

## Abstract

Model approaches to nuclear architecture have traditionally ignored the biophysical consequences of ATP-fueled active processes acting on chromatin. However, transcription-coupled activity is a source of stochastic forces that are substantially larger than the Brownian forces present at physiological temperatures. Here, we describe a first-principles approach to large-scale nuclear architecture in metazoans that incorporates cell-type-specific active processes. The model predicts the statistics of positional distributions, shapes and overlaps of each chromosome. Our simulations reproduce common organising principles underlying large-scale nuclear architecture across human cell nuclei in interphase. These include the differential positioning of euchromatin and heterochromatin, the territorial organisation of chromosomes including both gene-density-based and size-based chromosome radial positioning schemes, the non-random locations of chromosome territories and the shape statistics of individual chromosomes. We propose that the biophysical consequences of the distribution of transcriptional activity across chromosomes should be central to any chromosome positioning code.

## Highlights

- First-principles predictive model for large-scale nuclear architecture incorporating non-equilibrium activity
- Differential activity and looping patterns underly cell-type-specific features of such architecture
- Differential positioning of inactive and active X chromosomes an emergent property
- Simulations of the model recapitulate many known features of nuclear architecture and predict new ones

## Introduction

Chromosomes are not distributed at random within the interphase nucleus, an observation that is central to our current understanding of large-scale nuclear architecture in the interphase nuclei of metazoans (Meaburn and Misteli, 2007; Cremer and Cremer, 2010; Bickmore and van Steensel, 2013). Gene rich, more open, early-replicating euchromatin regions are typically distributed more centrally than gene-poor, relatively more compact, late-replicating heterochromatin (Cremer and Cremer, 2010). Chromosomes are organised territorially, with each being segmented into relatively more (A) and less (B) active compartments that are then further subdivided into topologically associated domains (Lieberman-Aiden et al., 2009; Dixon et al., 2012; Fraser et al., 2015). In humans, gene-rich chromosome 19, containing a large number of house-keeping genes, is distributed more centrally across several cell types than the similarly sized but gene-poor chromosome 18 (Croft et al., 1999; Boyle et al., 2001). This observation generalises to a gene-density-based radial positioning schema for all chromosomes (Takizawa et al., 2008). Gene-rich regions within chromosomes tend to orient towards the nuclear centre, with expressed alleles often found further from the nuclear envelope than ones that are not expressed (Takizawa et al., 2008; Therizols et al., 2014). In some human cell types, chromosomes appear to be positioned by size, with the centres of mass of smaller chromosomes disposed more centrally than those of larger ones (Sun et al., 2000; Bolzer et al., 2005; Kölbl et al., 2012). In female cells, the two X chromosomes are differentially positioned, with the more compact, inactive X-chromosome found somewhat closer to the nuclear envelope than the active one (Dyer et al., 1989; Jégu et al., 2017). Actively transcribed chromosomes tend to have rougher, more elliptical territories than less active ones (Eils et al., 1996; Berezney et al., 2005; Khalil et al., 2007; Sehgal et al., 2014; Jégu et al., 2017). The probability with which two loci along individual chromosomes are found in proximity to each other in ligation assays follows a power-law P(s) ~ 1/s^*α*^ with *α* ⋍ 1 over an approximately 1 - 8 Mb range, consistent with a fractal globule picture of chromosome structure (Lieberman-Aiden et al., 2009; Mirny, 2011). Currently, experiments suggest that such organization is cell-type dependent and that *α* (1 ≤ *α* ≤ 1.5) also varies across chromosomes over a comparable range (Sanborn et al., 2015; Kang et al., 2015).

Most model approaches to nuclear architecture assume *a priori* that chromosomes are structured polymers in thermal equilibrium (Cook and Marenduzzo, 2009; Tark-Dame et al., 2011; Marti-Renom and Mirny, 2011; Heermann et al., 2012; Vasquez and Bloom, 2014; Imakaev et al., 2015). Some models ignore thermal fluctuations altogether in favour of incorporating loop structure as derived from the Hi-C data, while also requiring compatibility with physical restrictions on the overlaps of chromosomes (Imakaev et al., 2015; Amitai and Holcman, 2017; Tjong et al., 2016). Others account for the domain structure of individual chromosomes (Odenheimer et al., 2005; Jost et al., 2014; Jost et al., 2017; Chiariello et al., 2015; Haddad et al., 2017; Ghosh and Jost, 2017; Zhang and Wolynes, 2017; Tiana et al., 2016; Di Pierro et al., 2016; Di Pierro et al., 2017). As summarized above, large-scale nuclear architecture exhibits generic features that are largely common across cell types. These should severely constrain potential models (Bickmore, 2013). However, set against this stringent requirement, virtually all prior models for such architecture are incomplete: (i) these models fail to predict gene-density based or size-based positioning schemes; (ii) no simulations reproduce the chromosome-specific distribution functions for gene density or chromosome centre-of-mass that FISH-based experiments provide; (iii) the differential positioning of the active and inactive X chromosomes cannot be obtained using any model proposed so far and (iv), the spatial separation of heterochromatin and euchromatin, seen in interphase cell nuclei across multiple cell types, has not been reproduced in model calculations in which this information is not incorporated *a priori*. Understanding these discrepancies remains an outstanding problem.

All molecular machinery associated with chromatin remodelling, transcription and DNA repair is energy-consuming, relying on the hydrolysis of NTP molecules (Flaus and Owen-Hughes, 2011). Recently, we pointed out that this leads to the localised, irreversible consumption of energy at the molecular scale (Ganai et al., 2014). This energy is transduced, through chemo-mechanical “active” processes, into mechanical work (Weber et al., 2012; Zidovska et al., 2013; Chu et al., 2017). Such processes can be modelled *via* recently developed biophysical theories of “active matter” (Menon, 2010; Prost et al., 2015; Marchetti et al., 2013; Needleman and Dogic, 2017). We argued that a description in terms of inhomogeneous, stochastic forces acting on chromatin, equivalent to an effective temperature reflecting local levels of activity, provided the right biophysical setting (Fodor et al., 2015; Hameed et al., 2012). Describing each chromosome as a polymer composed of consecutive monomers, each representing a suitably averaged section of chromatin, different monomers can then be expected to experience different effective temperatures correlating to local active processes (Ganai et al., 2014; Agrawal et al., 2017; Wang and Wolynes, 2011). Here, extending these ideas, we propose an *ab initio* biophysical approach to predicting both cell-type-specific and cell-type independent features of large-scale nuclear architecture, using data from RNA-seq experiments as a proxy for activity and a Hi-C-derived description of chromosome looping in each cell type. The model provides a unified understanding of a number of common features of large-scale nuclear architecture observed across diverse cell-types.

## Results and Discussion

We model human chromosomes in diploid female (XX) cells within interphase, describing each as a polymer made up of monomers linked along a chain. These polymers are confined within a spherical shell that models the nuclear envelope. Each monomer represents a 1Mb section of chromatin (Kölbl et al., 2012; Jackson and Pombo, 1998). Our model chromosomes are dynamic and explore different configurations, based on the forces they experience. Such forces arise from the dense, non-equilibrium and fluctuating environment of the cell nucleoplasm, the interactions of chromosomes and chromosome-nuclear envelope interactions. A number of simulation snapshots of both homologs of chromosomes 18 and 19, against a background of all other chromosomes represented in grey-scale, are shown in Figure 1A. From such snapshots, we compute a variety of statistical properties of chromosomes accessed in experiments (Figure 1B-G).

**Figure 1:**
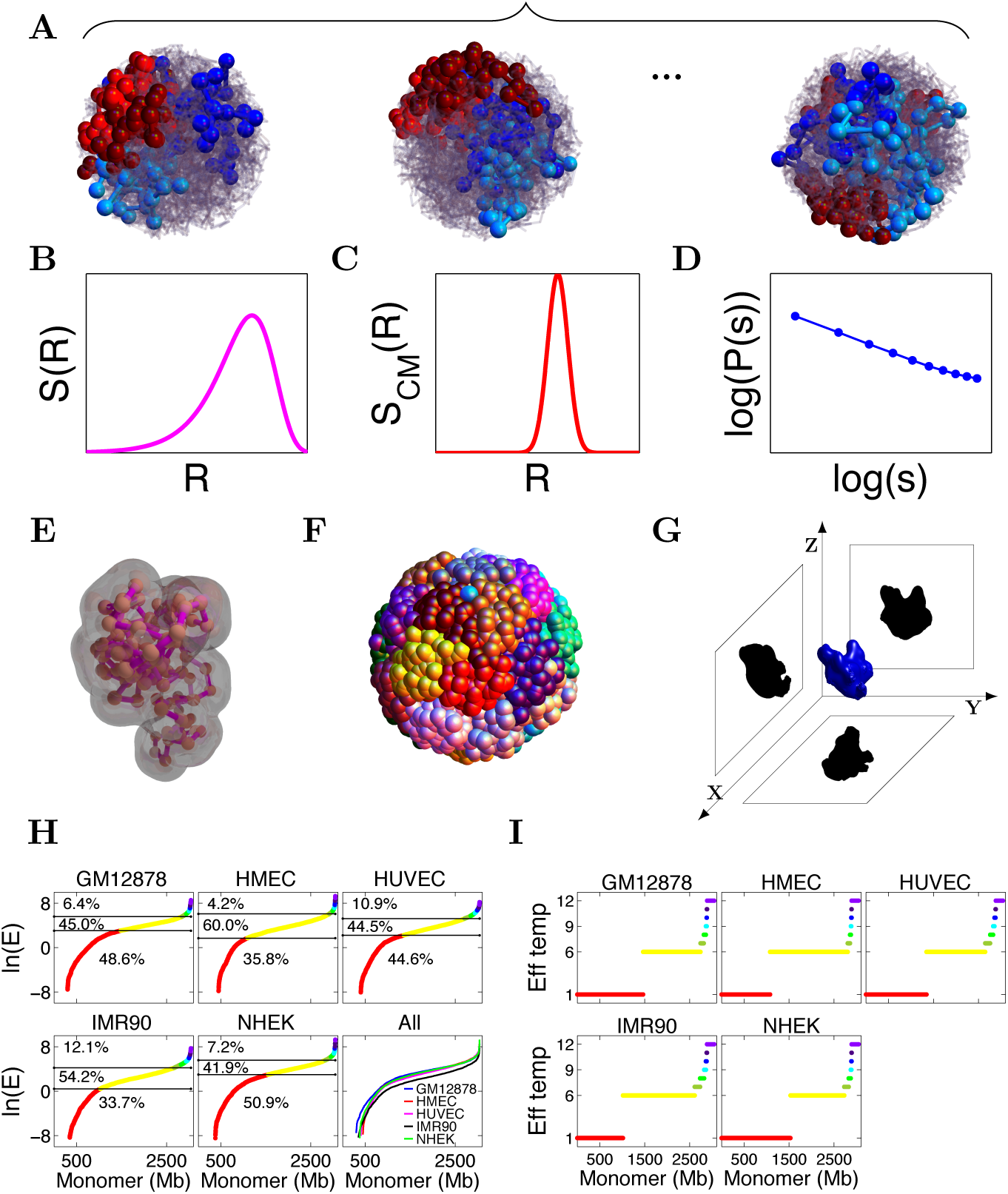
Model schematics and active temperature assignments. **A.** Several simulated configurations of 23 pairs of chromosomes within a spherical nucleus, with pairs of chromosomes 18 and 19 highlighted in the background of other chromosomes, shown in grey scale. Each bead represents a 1 Mb section on each chromosome. We average all calculated quantities, such as distribution functions, over a large number of such configurations in steady state. **B.** Schematic of the DNA distribution S(R) of each chromosome, plotted against the radial coordinate R and averaged over many nuclei in our simulations. **C.** Schematic of the centre of mass distribution of each chromosome, S_*CM*_(R), plotted against the radial coordinate R and computed from an average over many simulated nuclei. **D.** Schematic of the contact probability P(s) between beads of chromosomes, for two monomers separated by an internal (genomic) distance s along the polymer. **E.** The shapes of individual chromosome territories extracted from simulation configurations. Such shapes are used to compute a number of geometrical properties of chromosome territories, e.g. their volume, surface area, asphericity and other shape parameters. **F.** Typical image of chromosome territories computed in our simulations, with each chromosome colored a different color, illustrating the emergence of territoriality. **G.** Schematic illustrating a 2D projection of a three-dimensional chromosome territory, projected along the *XY*, *Y Z*, and *XZ* planes. The ellipticity and regularity parameters can be computed from such 2d projections, and compared to 2D FISH data. **H.** The logarithm of gene expression values for each 1Mb monomer, plotted in order of increasing gene expression. These are computed from transcriptome data. Data are shown for 5 cell types as indicated in the title to each sub-figure. The horizontal lines drawn motivate our assignment of effective temperatures as discussed in the text, and correspond to our assignment of activity in proportion to gene expression. The last sub-figure plots these data together, illustrating that the shape of the activity profile is largely similar, even though individual monomers in different cell types can be classified differently on the basis of their activity. **I.** Assignment of effective temperature to each monomer for the combined model. The red monomers are simulated at *T* = 1, yellow at *T* = 6, yellow-green at *T* = 7, green at *T* = 8, cyan at *T* = 9, blue at *T* = 10, indigo at *T* = 11 and violet at *T* = 12 times the physiological temperature T_*ph*_.

We work with three models that associate local levels of non-equilibrium transcriptional activity to an effective temperature. In the **gene density model**, proposed in our earlier work, we chose the top 5% of monomers by gene density, assigning them an active temperature in excess of the physiological temperature *T_ph_* (Ganai et al., 2014). The gene density model yields fairly accurate representations of the measured distribution function of DNA density S(R) in GM12878 cells, leading to very different distributions for the chromosome pairs 18 and 19 *vis a vis*. chromosomes 12 and 20, as seen experimentally (Ganai et al., 2014). However, such a model is insensitive to cell-type-dependent features of nuclear architecture (Bickmore, 2013). Accordingly, in the **gene expression model**, we focused on transcriptomes across a number of model systems, exploring varied ways of associating transcript levels to effective temperatures. Figure 1H shows RNA-seq derived FPKM values summed over 1Mb intervals, indexing transcript levels, across GM12878, HMEC, HUVEC, IMR90 and NHEK cell types. Their distribution follows a Gumbel form (Figure S1). The values in Figure 1H are plotted in increasing order of expression on a logarithmic scale. We chose structured effective temperature assignments that reflect the overall shape of this curve. Surprisingly, the gene expression model did not yield appreciably better results than the much simpler gene density model. To address this, we noted that transcript levels need not directly correlate to activity, since FPKM values are controlled by the rate at which transcripts are both produced and degraded, because non-coding transcription is not fully captured in this version of our model, and because our description averages over the typical time-scales associated with transcriptional “bursts” (Fraser and Bickmore, 2007; Chubb et al., 2006). We felt that a model which included features of both gene density and gene expression models should provide a more accurate representation of inhomogeneous cell-type-dependent activity (Murmann et al., 2005). Accordingly, we decided to assign monomers with a gene density above a present cutoff, the maximum active temperatures, as in the earlier gene-density model. All the results presented in this paper are for this **combined model**. Figure 1I shows temperature assignments, within the combined model, for 5 cell types. Such inhomogeneous (effective) temperature assignments, correlating both to gene density and transcription levels averaged over consecutive 1Mb sections of each chromosome, lie at the core of our work.

### Inhomogeneous activity underlies large-scale nuclear architecture

Figure 2A shows an example of simulation-derived chromosome territories for the GM12878 cell type, with each chromosome coloured a different colour. Simulations recover such territorial organization robustly, illustrating how territoriality is an emergent consequence of our model. Figure 2B shows a cut-away profile showing the averaged spatial distribution of active (white, *T/T_ph_ >* 1) and inactive (black, *T/T_ph_* = 1) monomers in the GM12878 cell type, extracted from snapshots of a typical configuration. Low gene density monomers, shown in black and representing heterochromatin, are enriched towards the boundaries of the nucleus, whereas high gene-density euchromatin regions, shown in white, preferentially occupy the bulk. In Figure 2C, we show a cutaway profile of the time-averaged effective temperature within our simulated nucleus, an indicator of local activity in each spherical shell centred around the origin. In Figure 2D we show the time-averaged gene density across spherical shells, in a similar visualisation. Gene densities, as well as activity, increase towards the nuclear centre.

**Figure 2:**
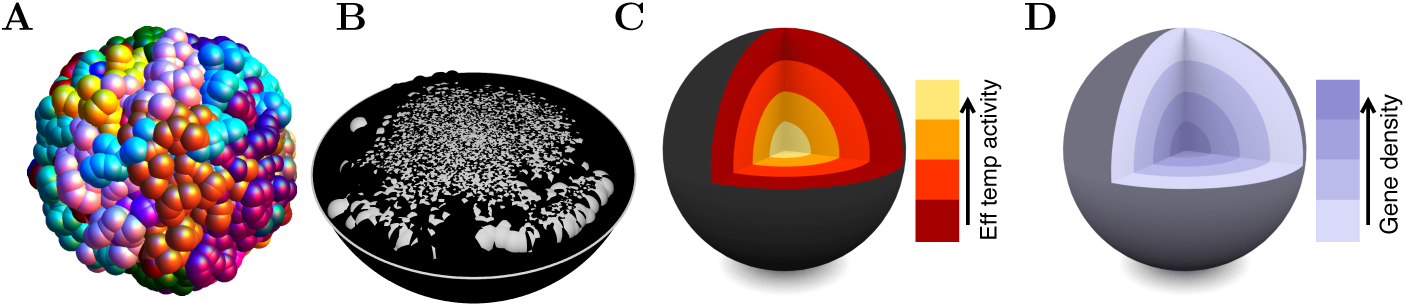
Model predictions for large-scale features of nuclear architecture. **A.** Chromosome territories computed in our simulations, with each chromosome colored a different color. Note the tendency of each chromosome to overlap relatively little, visually representing territoriality. **B.** A cut-away sphere representation of the average spatial distribution of euchromatin (or active white) and heterochromatin (or inactive black) monomers as computed for the GM12878 cell type. Here, the active monomers are defined as those having an effective temperature in excess of the physiological one. Heterochromatin is found more peripherally compared to euchromatin which is located towards the nuclear interior. **C.** A cutaway sphere representation of average effective temperatures within the simulated nucleus, as computed for the GM12878 cell type. This illustrates the larger effective temperatures, indicating enhanced activity, obtained towards the centre of the nucleus, in comparison to a lower effective temperature in the nuclear periphery. **D.** A cutaway sphere representation of the average gene density within the simulated nucleus, computed for the GM12878 cell type. This illustrates the excess in gene density seen towards the centre of the nucleus in comparison to the gene density in the nuclear periphery. This separation of gene-dense and gene-poor 1Mb segments of chromatin correlates to the distinction in the spatial positioning of euchromatin and heterochromatin.

Our *ab initio* biophysical description of chromosomes and their structuring reproduces the different spatial distributions of euchromatin and heterochromatin, a feature seen across multiple cell types. A central consequence of our model is that gene densities should correlate to a larger strength of mechanical fluctuations *i.e*. activity, and that the distribution of both these quantities should be attenuated towards the boundaries of the nucleus. This is an emergent property, arising from the combination of differential activity and confinement, that could not have been inferred from how the model was constructed.

### The model predicts positional distributions of individual chromosomes

Chromosome-specific distribution functions S(R) are obtained experimentally using confocal slices of FISH images from an ensemble of fixed nuclei. Our computed S(R) for chromosomes 18 and 19 in the 5 cell types we examine are shown in Figure 3A. All data are averaged over the two autosomal homologs, as their positioning was found to be equivalent. For the GM12878 cell type, we compare our results with experimental results extracted from Ref. (Kreth et al., 2004). S(R) for chromosomes 18 and 19 exhibit well separated peaks, a feature that holds across cell types. For comparison, the R^2^ rise of S(R) towards the nuclear envelope, expected for uniformly distributed chromosomes, is shown in each subfigure. Figure S2 shows our calculated S(R) for all chromosomes in the 5 cell types. Figure S3 shows S(R) for the GM18278 cell type where we compare the predictions of the gene expression model and the combined model. In Figure S4 we show how S(R) for the GM18278 cell type varies when we include or exclude looping and when we include or exclude activity. The predictions of the different models differ substantially for both the gene-rich chromosomes as well as the smallest chromosomes. In general, both looping and differential activity are needed to best represent available experimental data.

**Figure 3:**
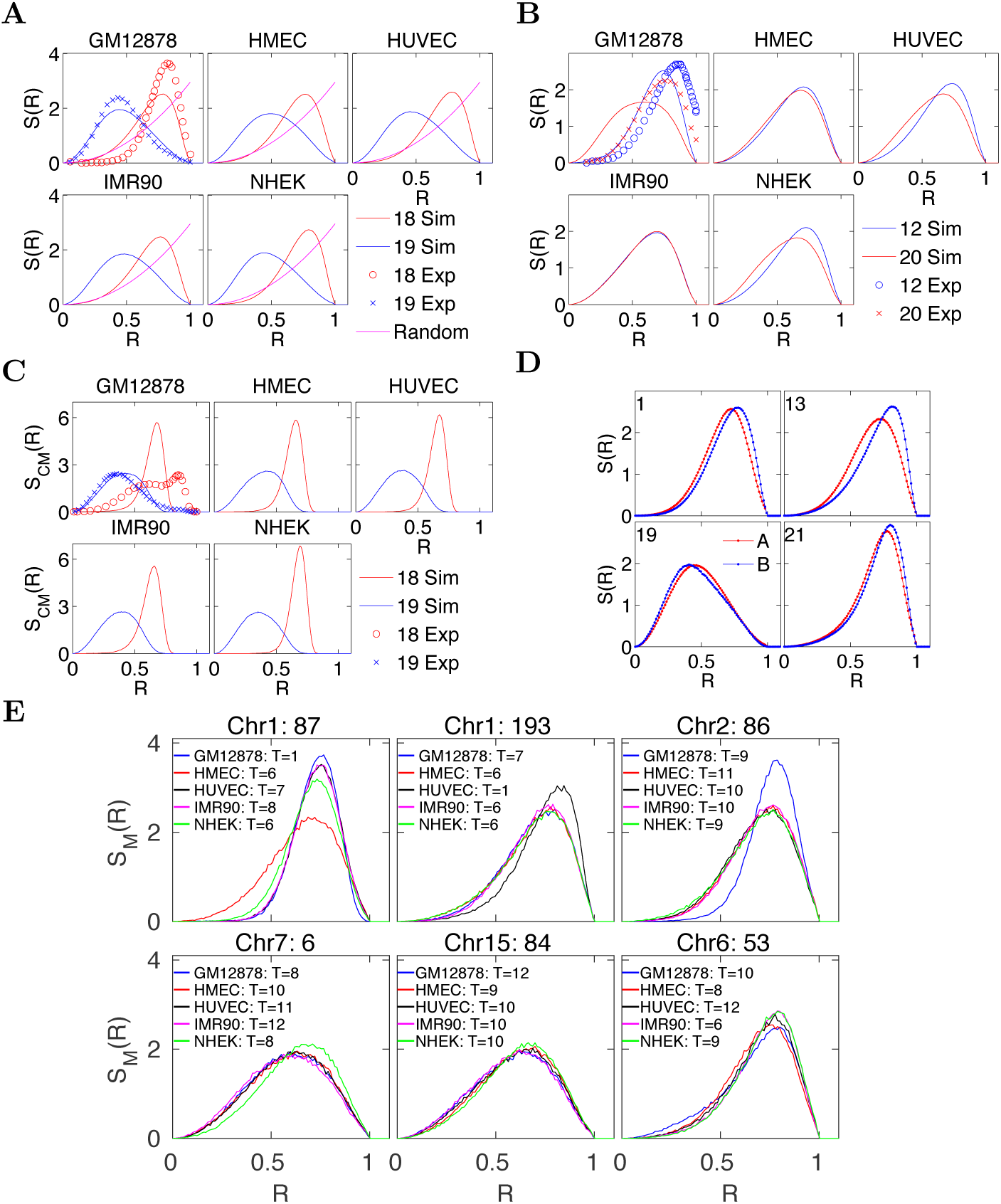
Predicted radial distribution functions S(R) compared to experimental data. **A.** Distribution of monomer density S(R), reflecting the local density of DNA, for chromosomes 18 and 19 (red and blue lines respectively) across 5 cell types as indicated in the titles of each sub-figure. Experimental data obtained from Ref. (Kreth et al., 2004) for the GM12878 cell type is plotted together with the simulation predictions, (red ovals: Chr 18, blue crosses: Chr 19). If chromosomes are distributed randomly across the nucleus, S(R) ~ R^2^ is expected, as shown with magenta lines. **B.** Distribution of the density of monomers, reflecting the local density of DNA, for Chr 12 and 20 (blue and red lines respectively) across 5 cell types as indicated in the titles of each sub-figure. Experimental data obtained from Ref (Kreth et al., 2004) for the GM12878 cell type is plotted (red ovals: Chr 18, blue crosses: Chr 19), together with the simulation prediction. **C**. Distribution of chromosome centres of mass for Chr 18 and 19 (red and blue lines respectively) shown for 5 cell types, as indicated in the titles of each sub-figure. Experimental data obtained from Ref. (Kalhor et al., 2011) for the GM12878 cell type are plotted (red ovals: Chr 18, blue crosses: Chr 19) together with the simulation prediction. **D.** Density distribution S(R) of overall numbers of active (red) and inactive (blue) monomers for the GM12878 cell type. These are plotted for 4 chromosomes largest Chr1, smallest Chr 21, gene poor Chr 13 and gene rich Chr 19. The distribution of active monomers is more interior with respect to inactive monomers. Here, inactive monomers refer to those monomers assigned a temperature of T = 1; all other monomers are active. **E.** Density distribution S_*M*_(R) of specific monomers as indicated, on chromosomes 1, 2, 7, 15, and 6, plotted for 5 cell types studied here. These monomer-specific distributions can differ depending on cell type, suggesting that loci associated to these monomers can be positioned differently depending on their levels of activity, but also on the levels of inhomogeneous activity of the chromosome they belong to.

Figure 3B shows S(R) for chromosomes 12 and 20. For the GM12878 cell type, we compare our results with results from Ref. (Kreth et al., 2004). Note that simulation data for the different cell types all yield similar plots for S(R), with the exception of the GM12878 cell type where, although the simulation and experimental data peak at different locations, the overall shape of the curve is rendered accurately, including the relative shift in peak positions. Figure 3C shows the distribution of centres of mass of specified chromosomes, S_*CM*_(R), for chromosomes 18 and 19. For GM12878 cells, we compare our results with experimental data from Ref. (Kalhor et al., 2011). The centre of mass distributions are captured well, especially for chromosome 19. The broader distribution of S_*CM*_(R) for chromosome 18 is also in agreement with the left tail of the experimental data, although the experiments show a weaker and more outward shifted peak than the simulation prediction. Differences in positioning of Chromosomes across cell types are more apparent in S_*CM*_(R) compared to S(R). Figure S5 shows _*cm*_(R) for all cell types across all chromosomes. We compare the predictions for S_*CM*_(R) in the gene expression and combined models in Figure S6. Results for S_*CM*_(R) with different combinations of activity and loops are shown in Figure S7. Overall, apart from the gene rich chromosomes, the predictions of the gene expression and combined models are comparable. The largest variability across cell types is seen in chromosomes 1, 4, 7, 11, 12, 16, 21 and 22. S_*CM*_(R) for gene poor chromosomes are sharply peaked while gene rich chromosomes have broader distributions across all cell types.

Figure 3D shows the partial distribution functions S(R) for inactive and active monomers in the GM12878 cell type for chromosomes 1, 13, 19 and 21. The distribution for active monomers is shifted towards the nuclear centre whereas for the inactive monomers, it is seen to be shifted towards the nuclear periphery. These results relate to the experimental observation that active alleles are positioned more towards the interior of the nucleus, an effect strong enough to be apparent in our simulations (Fedorova and Zink, 2009; Therizols et al., 2014). In Fig. 3E we show monomer-specific distribution functions S_*M*_(R) for 6 monomers across Chr 1, 2, 6, 7 and 15, across all cell types; these monomers contain multiple genetic loci and typically show differential activity across the cell types shown. We note that such monomer distributions are not identical but depend on both their active temperature as well as the overall activity and loop content of the chromosomes that contain them.

These results indicate that cell-type dependent signatures of activity can be especially prominent at the level of individual monomers, and thus loci. They are overall less prominent in chromosome-specific DNA density distributions and the distributions of their centres of mass, but display subtle differences nevertheless. These differences originate both in differences in activity profiles across different cell types as well as variations in their loop content, suggesting that these should be essential components of any biophysical description of large-scale nuclear architecture.

### Model predictions for size- and gene-density-dependent chromosome positioning

Figures 4 shows our computation of the mean centre of mass of each chromosome within the combined model for the GM12878 cell type, as compared to an analysis of three sets of experimental data for this cell type. These data, on the average radial position of each chromosome, are obtained through (A) FISH, (B) in-situ modeling of Hi-C data and (C) from the TCC (tethered chromosome conformation capture) data, all extracted from Fig S8 of Ref. (Tjong et al., 2016). Also shown are fits to straight lines as a function of chromosome size and as a function of gene density, independently for both experimental and simulation data. Simulation and experimental points are shown using red and blue filled circles respectively, together with error-bars indicating one standard deviation from the mean. The fit against chromosome size for the simulation data excludes the two smallest (21 and 22) chromosomes in the upper row (subfigures 1A-1C). In the lower row subfigures 2A-2C, all chromosomes are included in the fit. A measure of the quality of the fit, the Pearson correlation coefficient (PCC) *r*, is provided for each subfigure.

**Figure 4:**
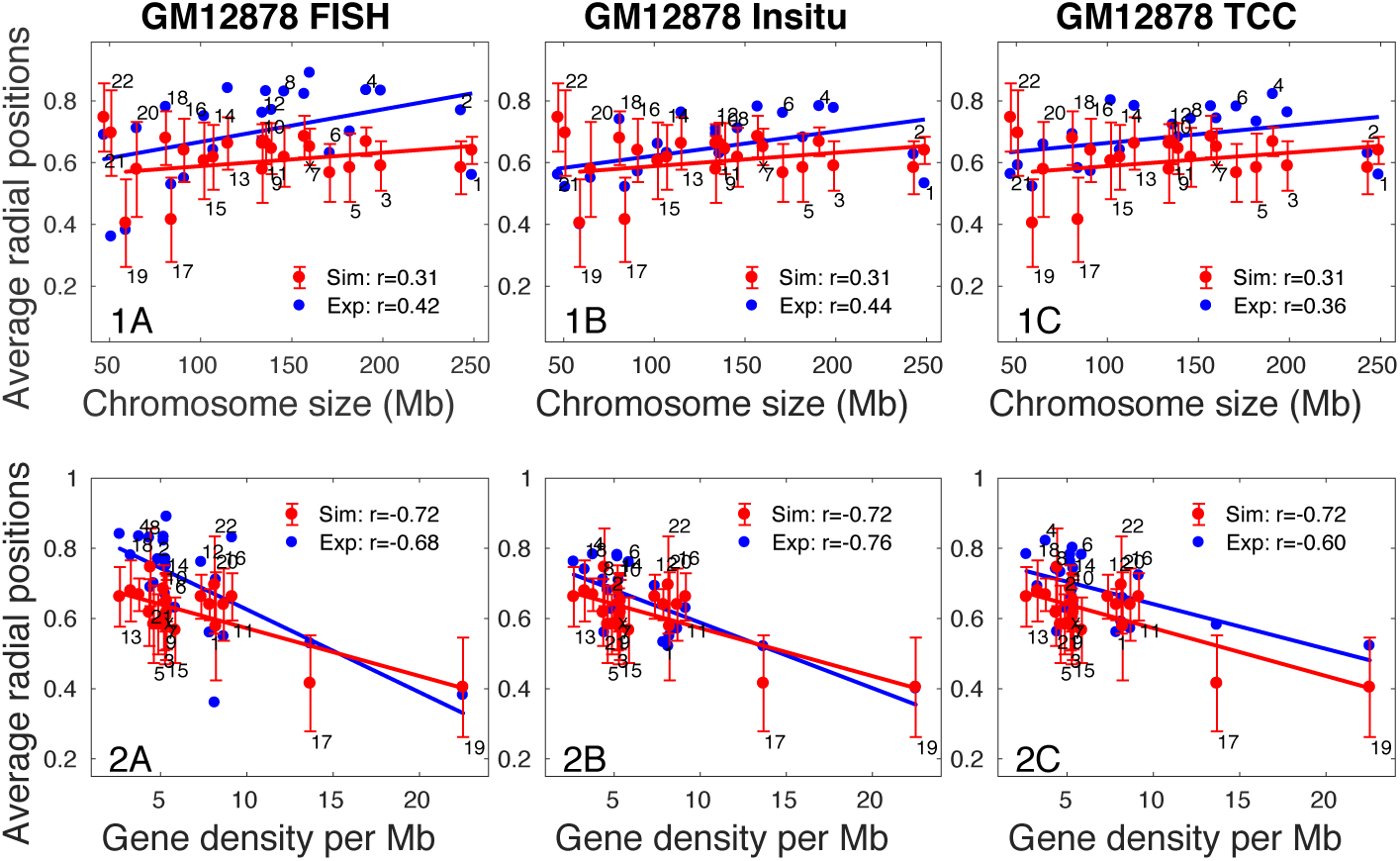
Predicted chromosome centre-of-mass locations compared to experimental data. Predictions from simulations for the mean centre-of-mass location for each chromosome, for the GM12878 cell type, as a function of chromosome size in the upper row (1A - 1C) and as a function of chromosome gene density per Mb in the lower row (2A - 2C). These predictions are compared to experimental data on the average radial position of the centre of mass of each chromosome as obtained through (1A, 2A) FISH, (1B, 2B) through in-situ modeling of Hi-C data (1B, 2B) and (1C, 2C) from the TCC data. All date has been extracted from Fig S8 of Ref. (Tjong et al., 2016). Simulation and experimental points are shown using red and blue filled circles respectively, together with errorbars indicating one standard deviation from the mean. The relative radial position 0 and 1 represent the centre and periphery of the nucleus. Chromosome numbers are indicated above or below each error bar. The simulation points, excluding the two smallest chromosomes 21 and 22, are fit to a straight line (see text), while the fit to the experimental points includes all chromosomes. The Pearson correlation coefficient (PCC) r between chromosome size and average radial positions, both for experiments and simulations, is provided in each subfigure.

The simulations reproduce most of the experimental systematics. The positions of experimental and simulation data coincide for some chromosomes or lie within error-bars. The positions of chromosomes 14, 16, 18, 19, 20 and X are very close to the experimental data, reproducing the unusual non-monotonicity in their positions. The slight overall shift between the positions between experiment and simulation arises from the fact that our simulations are performed for spherical nuclei whereas the experiments are performed on more flattened, ellipsoidal nuclei as well as averaged over an ensemble of such shapes.

The fits to the data are consistent with an approximate size-dependence of chromosome positions relative to the nuclear centre, although with a PCC value of simulation (Sim) *r* ~ 0.31, compared to experimental (Exp) numbers in the range 0.36 ≤ *r* ≤ 0.44. These are indicative of a relatively weaker correlation between chromosome centre-of-mass and relative positioning within the nucleus, when compared to the correlation between such positioning and chromosome gene density (see below). The activity associated with each individual chromosome also plays a role in determining its position. The mean centre-of-mass locations for chromosomes in different cell types are similar but not identical. Chromosomes 18 and 19, although similarly sized, have very different positions relative to the nuclear centre, as also seen in the data of Figures 3A and 3C. Note that chromosomes 21 and 22 in Figure 4A are positioned more towards the exterior of the nucleus in the simulations than in the experimental data.

When chromosome centres-of-mass are plotted against individual gene densities the slope of the straight line is negative in all cell types (Figures 4A and S8). In addition the PCC values computed both for the simulation data (Sim) as well as the experimental data (Exp) is larger than was the case for the chromosome size data (Sim: *r* ~ 0.72, Exp: 0.60 ≤ *r* ≤ 0.76. Thus, depending on the region that is fit, one can have fits to both size dependence and gene density dependence of chromosome centres of mass relative to the nuclear centre, but our methods predict that the gene-density dependence is the stronger one. The fact that the smallest chromosomes, Chr 21 and 22, lie outside of the fit to chromosome size may reflect aspects of their activity that our method does not resolve, as well as variations in loop assignments. Figure S8 show similar plots for simulation calculations of chromosome centres-of-mass plotted for HMEC, HUVEC, IMR90 and NHEK cell types.

Figure S9 shows the mean centre of mass position as computed for the GM12878 cell type, across a variety of simulation conditions, including for the gene density model as well as for the combined model with various choices for the incorporation of loops and activity. Figure S9A shows results for the gene expression model. In Figure S9B we show results for the case in which we allow differential activity but ignore looping. In Figure S9C we show results for the case in which differential activity is absent but looping, as prescribed by the Hi-C data, is retained. All monomers then experience the same effective temperature, which we take to be the thermodynamic temperature. Finally, in Figure S9Ds, we show results for the case where looping is absent as well, so that this case corresponds to the case of chromosomes without loops at thermal equilibrium. From these, we conclude that in the absence of both activity and looping, chromosome positioning is only weakly structured. Our simulations indicate that chromosome positioning is very weakly size-dependent or even independent of size in all conditions where activity is switched off. Allowing for loops induces some changes in positioning but these results do not match with experiment. Allowing for activity, but ignoring loops, leads to a differential positioning of chromosomes but, if anything, the size dependence of chromosome positions is *opposite* to that seen in the data. Only models which incorporate both activity and looping are successful in reproducing both the approximate size dependence of radial positioning, while also accounting for those specific cases which fall outside this general trend, such as chromosome 19.

The model predicts the centre-of-mass positions of most chromosomes with reasonable accuracy, well within the error bars on the measurements for virtually all chromosomes. Finally, the fact that a number of broad features of the experiments are reproduced in the model suggests that the large-scale structure and positioning of individual chromosomes are principally determined by inhomogeneous activity across chromosomes, the presence of loops and confinement.

### Shapes and statistical features of individual chromosome territories compare well to experiments

Figures 5A and 5B show territories for the two autosomes corresponding to chromosomes 12, 20, 18 and 19. In Figure 5C, we show comparisons between 2d FISH data for chromosome regularity and ellipticity on WI38 cells, for which data is available (Sehgal et al., 2014), to predictions from our simulations for the GM12878 and IMR90 cell types. Both IMR90 and WI387 are lung fibroblast cell lines. Chromosomes are indexed, along the x-axis, in order of their gene density. The simulations and experimental data track each other, with the simulations finding the same dip, and subsequent rise, of both ellipticity and regularity around chromosome 22. Both ellipticity and regularity peak for chromosome 11, a feature both of the simulations and of the experiments. The ellipticity and regularity also appear to decrease weakly with increasing gene density, although individual chromosomes may deviate from this general trend (Figure 5C).

**Figure 5:**
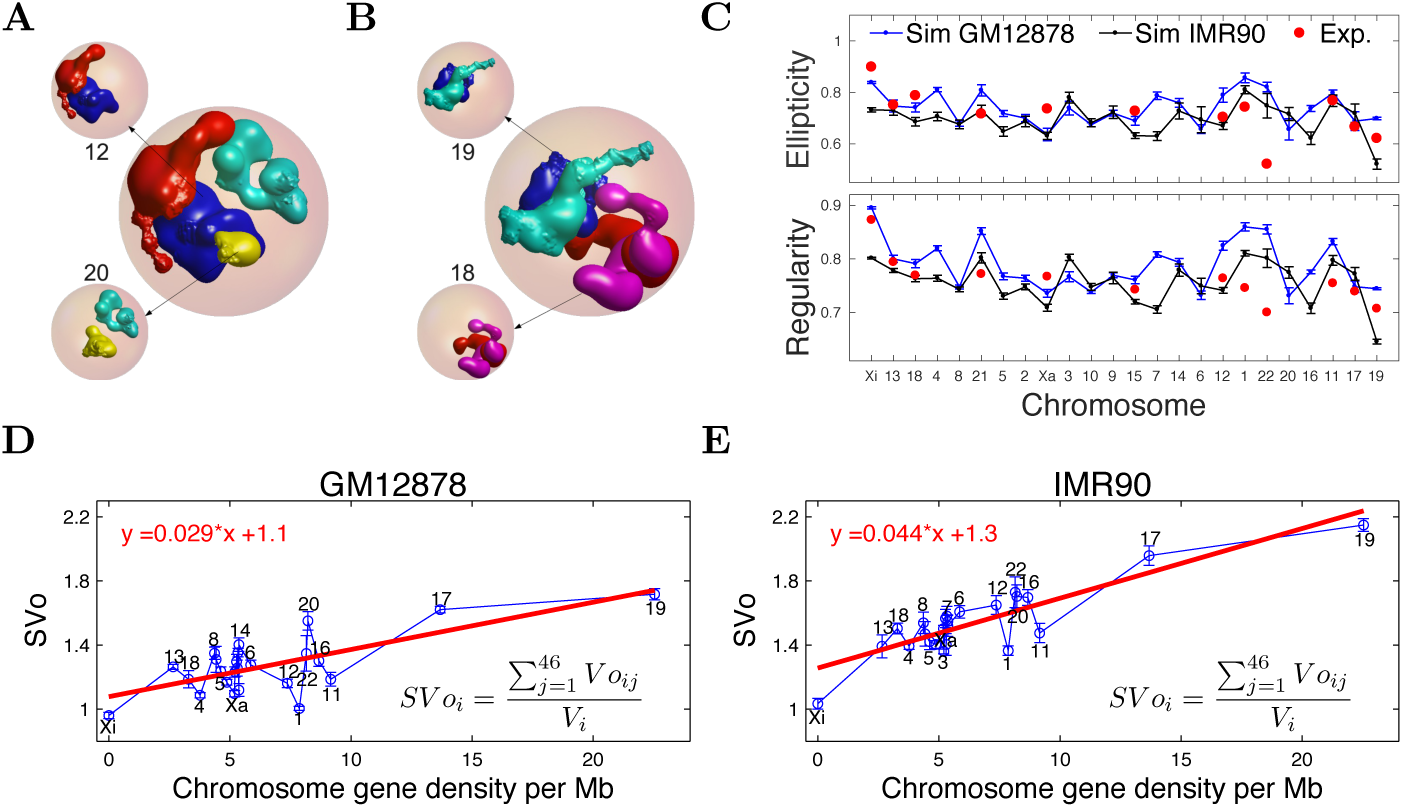
Structural properties of individual simulated chromosomes in our model. **A.** Snapshot of simulated configurations of both homologs of chromosomes 12 and 20. Each chromosome is colored differently so that they can be separately visualized. **B.** Snapshot of simulated configurations of both homologs of chromosomes 18 and 19. Each chromosome is colored differently so that they can be separately visualized. **C.** Ellipticity and Regularity for each chromosome as predicted by the model and obtained from simulations representing the GM12878 (blue) and IMR90(black) cell types. These are compared to experimental data (red oval symbols) from 2d FISH experiments Ref. (Sehgal et al., 2014) for a cell type closely related to the IMR90 cell type. Ellipticity values of 1 represent a perfect elliptical chromosome and regularity values of 1 refer to a perfectly regular chromosome, without roughness. The x axis is plotted in order of increasing gene density. **D-E**. Summed volume overlap (SVo) of chromosomes in GM12878 and IMR90 cell types, with the x-axis plotted in order of increasing gene density per chromosome. There is a weak increase with gene density in both cell types, shown as the solid line, representing the best linear fit to the data. The IMR90 cell shares more volume overlaps with other chromosomes compared to the GM12878 cell type. The (self-) volume overlap (Vo) for the same chromosome is taken to be 0.

Figures 5D and 5E show the summed volume overlap (SVo), sometimes referred to as the intermingling, and used to understand chromosome-chromosome interactions in trans, of different chromosomes in our model. The ordering of chromosomes according to their gene density per chromosome as shown on the x-axis is the same as the ordering used for the 2d projected data in Figure 5C. The largest overlap is for the most gene rich chromosome. There are perceptible differences in the overlaps of chromosomes in the GM12878 and the IMR90 cell types.

In summary, the simulations reproduce broad features of individual chromosome territories. More active chromosomes deviate more from a spherical shape and have rougher territories (Berezney et al., 2005). The summed volume overlap increases approximately linearly with chromosome gene density, with the Xi being an exception to this trend. Activity and looping have countervailing trends, since activity expands chromosome territories while looping contracts them.

### Simulations reproduce the differential positioning of the active and inactive X chromosome

Experiments investigating the positioning of the active and inactive homologs of the X chromosomes within interphase have consistently found that they are differentially positioned. The inactive X chromosome (the heterochromatic Barr body) is located most often towards the periphery of the nucleus (Jégu et al., 2017). This contrasts to the more central disposition of the active X chromosome Xa, which is larger and more extensively transcribed than the more compact Xi. We thus specifically investigated the positioning and other structure properties of the Xa and Xi chromosomes, since we expected that they would provide an example of where our methods, which emphasise the role and importance of activity, would yield predictions that other models could not.

Figure 6A shows a simulation snapshot of active and inactive X chromosome territories. Figure 6B shows our predictions for how these chromosomes are differentially positioned across all the cell types we study, through S(R). For the GM12878 cell type, we calculate S(R) for the Xi in two ways from simulations: In the first, shown with a red dashed line, we ignore the presence of “superloops”. These are recently studied large-scale loops that provide additional compaction in inactive X chromosome. In the second, shown with a red solid line, our simulations account for such superloops. The inactive X chromosome (Xi) has an S(R) which is sharply peaked close to the nuclear periphery. Accounting for superloops leads to a narrower S(R) distribution. Although the active X chromosome has a peak at a comparable location, its distribution has a long tail towards the nuclear centre. Figure 6C shows the calculated distribution of the centre of mass S_*CM*_(*R*) for these multiple cell types, verifying this essential distinction. Here again, for the GM12878 cell type the red dashed line is the case without superloops, while the red solid line is for the simulations that include them. This distinction between the distribution of Xa (blue solid line) and Xi (red solid line) in Figs. 6B-C suggests that differential positioning should be more readily seen in S_*CM*_(*R*) than in S(R).

**Figure 6:**
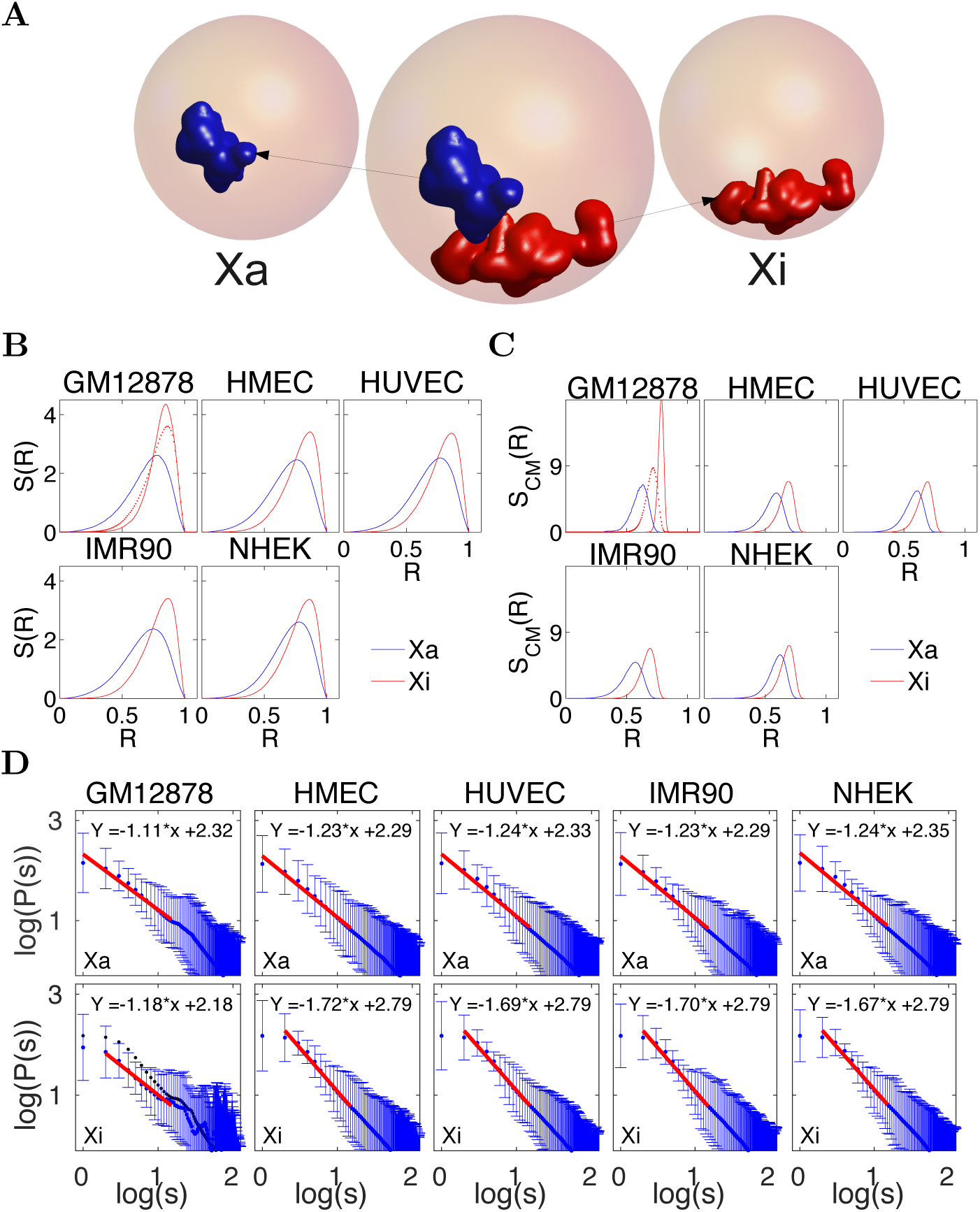
Predicted distributions and structural features of the inactive and active X chromosomes. **A.** Snapshot of a typical configuration of the active Xa and inactive Xi chromosome obtained from simulations. **B** Monomer density distribution S(R) vs R, for the Xi and Xa chromosome as obtained from simulations across 5 cell types, named in the header to each subfigure. The inactive X chromosome, Xi, is shown in red and the active X chromosome, Xa, is shown in blue. Loops on the Xi in the GM12878 cell type can include (red solid line) or exclude (red dashed line) “superloops” as seen in recent experiments Ref. (Rao et al., 2014; Darrow et al., 2016) **C.** Distribution of the location of the centre-of-mass of the Xi and Xa chromosome as obtained from simulations across 5 cell types, named in the header to each subfigure. The inactive X chromosome, Xi, is shown in red and the active X chromosome, Xa, is shown in blue. Superloops on the Xi in the GM12878 cell type can include (red solid line) or exclude (red dashed line). **D.** Contact probability P(s) vs s, for the active (top row) and inactive (bottom row) X chromosomes, computed for 5 cell types within our simulations. The Xa chromosome exhibits a reasonable power-law decay of P(s) with an exponent *α* between 1.1 and 1.25. The Xi chromosome shows a reduced region of power-law scaling, with an exponent across this reduced range which is between 1.5 and 1.7. Red lines show the power-law fit in both cases, with the fit parameters indicated within each sub-figure. In the absence of superloops on the Xi (GM12878) *α* ⋍ 1.5 (fitted line not shown) while the fit in the presence of superloops reduces *α* to α ⋍ 1.18.

We can compute the contact probabilities P(s) by applying a cutoff to the monomer-monomer distance distributions obtained in our simulation, averaging across a large number of simulation configurations. Figure 6D shows our computation of the contact probability P(s) for both Xa and Xi, across the 5 cell types we study. The cell type is shown at the top of each sub-figure. The active X chromosome shows more prominent power-law scaling of the contact probability than the inactive X chromosome, where any fit to a power law can only be over a far shorter genomic scale. Exponents for the power-law scaling of P(s) range from 1.11 - 1.24, with the smallest values obtained for the GM12878 cell type. For the Xi chromosome, accounting for superloops leads to a comparable scaling. However, in other cell types, such superloop information is unavailable. Accounting for loops as obtained through conventional Hi-C leads to a power-law exponent obtained varying from 1.52 - 1.72 in those cell types. The variation in the scaling of P(s) between Xa and Xi should be accessible experimentally, but the presence of superloops in other cell types as well might lead to a smaller divergence between these cases.

### Structural features of individual chromosomes are well described in our model

Figure 7A exhibits our results for contact probability P(s) of chromosome 1, across the five different cell types we study here. The data for small s show a power-law P(s) ~ 1/s^*α*^ behaviour over approximately a decade. The exponent is smallest for the GM12878 cell type, where our fits yield *α* ≈ 1.06. This value is very close to that obtained experimentally across the same region of genomic separation (Lieberman-Aiden et al., 2009; Sanborn et al., 2015). Values of *α* for all other cell types are consistently larger, with the exception of the IMR90 cell type. Overall, fitting *α* directly to the data across cell types yields 0.97 ≤ *α* ≤ 1.27. We see *P*(*s*) ~ 1/s^*α*^ with *α* ⋍ 1 over a 1 – 10 Mb range, as predicted by the fractal globule model, even though our model lacks virtually all the requisite ingredients for this model to be applicable. All we require is that activity is differentially distributed along the chromosome, that we account for looping as drawn from the Hi-C data, and that we account for crowding by other chromosomes, all features that previous work elides. Figures S10 and S11 shows plots of P(s) for all chromosomes computed for the GM12878 and IMR90 cell type. This P(s), for each chromosome, is best described within the 1 - 10 Mb range in terms of a wide range of exponents in the range 0.97 ≤ *α* ≤ 1.40.

**Figure 7:**
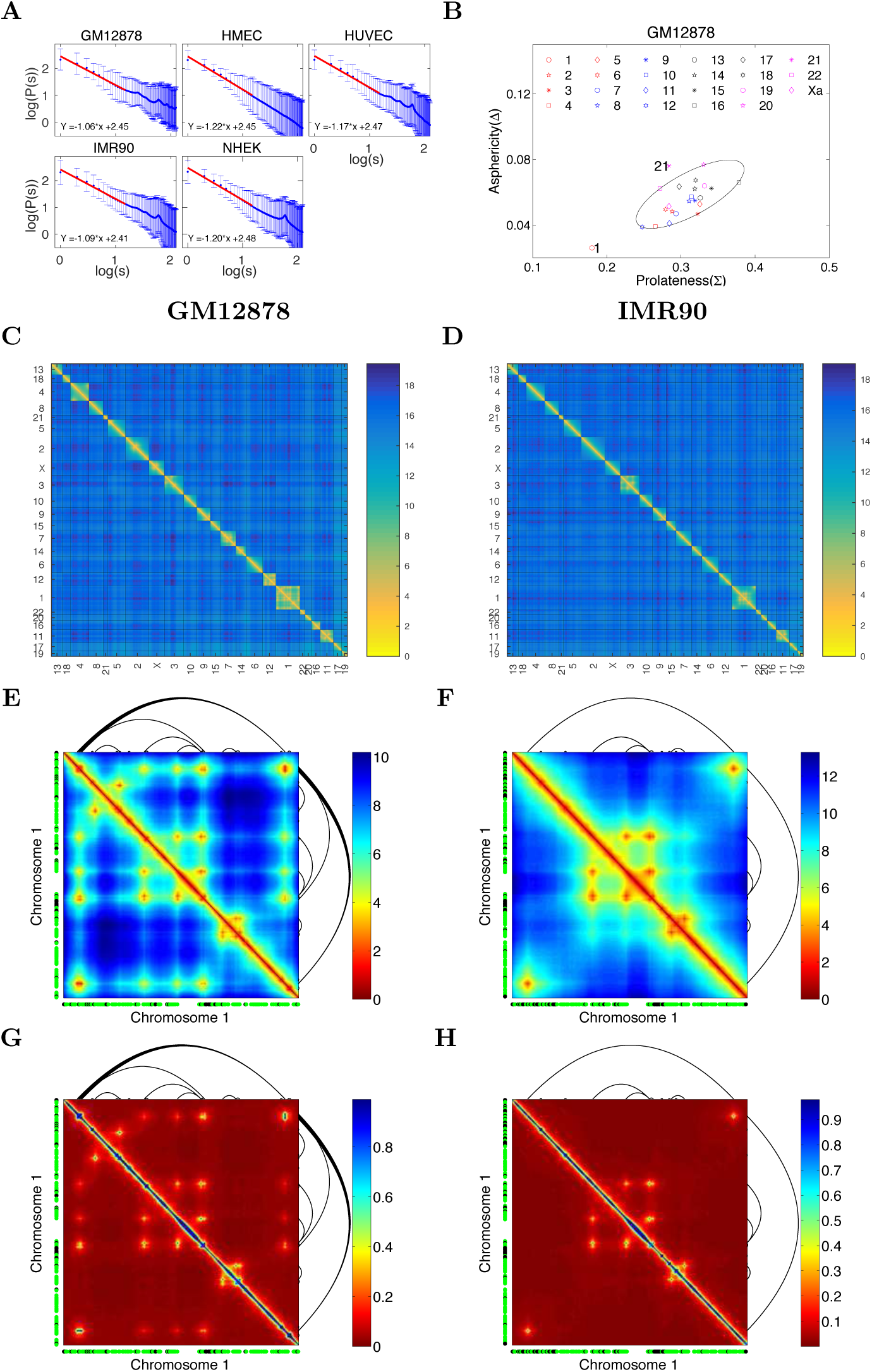
Predicted chromosome contact probabilities, distance maps and contact maps. **A.** Contact probability P(s) as a function of genomic distance for Chr 1, computed across a range of 1 - 15 MB and plotted for 5 different cell types. Our data is plotted with blue dots displayed with error-bars. Depending on the region that is fit, a power law scaling is obtained with an exponent between roughly 1.17 and 1.22; these fits are shown with red colors. **B.** Calculated average values of the prolateness parameter (Σ) and the asphericity parameter (Δ) for the GM12878 cell type. Larger (smaller) chromosomes have smaller(larger) values of Σ and Δ, implying that larger chromosomes are more close to spherical, while smaller chromosomes prefer a more prolate, rod-like shape. The data suggests that values of Σ and Δ for Chr 1 and 21take more extremal values than for the other chromosomes, as shown by the ellipse drawn together with the data. **C.** Heatmap of mean distances between monomers, the distance map, in which chromosomes are ordered by their gene density, shown for the GM12878 cell type. **D.** Heatmap of mean distances between monomers, the distance map, in which chromosomes are ordered by their gene density, shown for the IMR90 cell type. **E-F.** Heatmap of the distance matrix for chromosome 1, expanded out from Figures 7C and 7D. The locations of the permanent loops inferred from the Hi-C data are plotted in black. Individual monomers at T = 6 and 7 ≤ T ≤ 12 are shown in green and black, adjacent to the X and Y axis, respectively. **G-H.** Contact map inferred from the distance matrix for chromosome 1, (Figures 7C and 7D). The locations of the permanent loops inferred from the Hi-C data are plotted in black. Individual monomers at T = 6 and 7 ≤ T ≤ 12 are shown in green and black, adjacent to the X and Y axis, respectively.

Our model specification can be relaxed in several ways, so that we can examine and quantify independent contributions to this behaviour. For the specific case of chromosome 1, we have also investigated the predictions of the gene expression model, as shown in Figure S12A, as well as the effects of varying both activity and looping in the combined model (Figures S12B-S12D). Both the gene expression and combined models exhibit values of *α* which lie close to the experimental data, for which *α* ⋍ 1. In Figures S12B-S12D, we show results for the combined model with varying combinations of activity and looping. In the absence of both activity and loops, the exponent is close to the *α* = 1.5 expected for simple polymers. Adding loops or activity reduces this exponent. However, only the combined model, which includes both activity and looping obtains *α* values closest to those in experiments.

In Figure 7B, we show the spread of the asphericity parameter Δ and the shape parameter Σ, across chromosomes in the GM12878 cell type. (We exclude Xi, since its behaviour appears to depend sensitively on whether super-loops are included.) The simulations yield a linear relationship between Δ and Σ. Larger chromosomes have smaller values of Δ and Σ. Thus, an outcome of our model is that larger chromosomes are more spherical. The regularity and ellipticity indices calculated for the 2-d projections are in reasonable agreement with experimental trends (Figure 5C). We find that the data appears to fall into two classes, one a more compact set corresponding to all chromosomes with the exception of Chr 1 and 21 contained within an elliptical domain as shown in Figure 7B. Values of Δ and Σ for these special three chromosomes appear to be somewhat displaced from the locations for the other chromosomes, falling approximately onto the periphery of a larger ellipse. We show similar plots for other cell types in Figure S13. In the absence of activity, both if loops are present or absent, the Δ and Σ values for these chromosomes falls within the inner elliptical region across the 5 cell types we consider (Figure S14).

Figure 7C shows a heat map of monomer distances of chromosomes, indexed in increasing order of gene density for the GM12878 cell type. A similar plot is shown for the IMR90 cell type in Figure 7D. One feature of the data is that the more active chromosomes show smaller values of inter-chromosomal distance, likely reflecting the fact that more active regions are enriched towards the nuclear centre. In Figures 7E and 7F, we show the enlarged distance maps for chromosome 1. Applying a cutoff to such data, we can derive the likelihood of contacts arising from intra-chromosomal interactions, yielding P(s). Solid lines outside the figure body indicate those permanent attachments between different monomers that the Hi-C data provides. Note that regions connected by such loops exhibit a larger overlap. Figures 7G and 7H shows the contact maps inferred after applying a cutoff to the corresponding distance map. The borders of the axes show, in black and green, the active temperatures associated to specific monomers belonging to those chromosomes. The black colour refers to the most active monomers, with an effective temperature of 12 in units of the physiological temperatures whereas the green colour shows monomers with an effective temperature in the range 6 – 11. Monomers with a lower effective temperature are not shown. Regions with the same high effective temperature appear to contact each other more, but these are further modulated by the presence of internal loops. Note the presence of a dark banded region towards the centre of chromosome 1, associated to a large inactive region on this chromosome. This is a prominent feature of the experimental data, also seen in other cell types (Rao et al., 2014). We display similar plots for other cell types in Figure S15.

To summarise, our model yields structural information for chromosome structures and shapes that are broadly in agreement with available data. Our simulated distance maps lack the fine detail of distance maps computed in Hi-C experiments, which provide data for contacts at the smaller scales of 10 - 100 kB, but nevertheless are relevant to experiments that probe large-scale structuring. Our computed P(s) contains about a decade or so of power-law decays, with exponents that are comparable to those seen in experiments. Our model-based predictions for trends in the asphericity and prolateness of chromosomes with chromosome size and gene density are testable.

## Conclusions

Model descriptions of chromosomes must bridge multiple scales, ranging from microscopic length-scales of a few angstroms to scales of microns, of order the nuclear size. For now, brute-force atomistic simulations of the 23 pairs of chromosomes in human nuclei contained within the densely crowded, fluid and confined environment of the nucleoplasm are impossible. They are likely to remain so at least for the foreseeable future. Understanding how microscopic descriptions connect to macroscopic ones thus requires intuition for the processes that act to couple these scales, so that model building, which is as much about what to leave out as it is about what to leave in, can proceed.

The model described in this paper stresses a specific biophysical effect, ignored in previous work, of relevance to the modelling of chromosomes in living cells. We began by emphasising the relevance of non-equilibrium effects arising from local transcriptional activity for descriptions of nuclear architecture (Chu et al., 2017; Almas-salha et al., 2017). We proposed that the intensity of active processes should increase with increased transcription levels. We mapped a reasonable measure of local transcriptional activity, inferred from combining population-level measures of local RNA-output with estimates of the local gene density, into an effective temperature seen by each monomeric unit in our polymer model of chromosomes. We then performed simulations of these confined polymers, with properties chosen to reflect generic biophysical aspects of chromosomes. The monomers in our simulation represented 1Mb sections of chromosomes, although we could have defined our model at the smaller scales of 0.1 or even 0.01 Mb. However, the averaging inherent in summing transcriptional output over a 1Mb scale renders the model relatively less sensitive to errors and noise in this input. Further, the 1Mb scale is believed to be an appropriate building block for chromosome territories. A more detailed and explicit model for non-equilibrium activity and its consequences for an active temperature description would be useful, but the form such a model ought to take is presently unclear and best left to more extensive investigations. Irrespective of potential quantitative improvements on the model front, the broad trends we describe here should be largely robust.

We made a number of other modelling choices. First, we ignored the important role of lamin proteins in anchoring specific lamin-associated domains (LADS) to the nuclear lamina, as well as the interactions of specific gene loci with nuclear pore complexes (Mattout et al., 2015). While these are important omissions, they can at least be qualitatively justified by the biophysical intuition that the activity-based physical segregation of chromosomes is a bulk or “volume” effect that should dominate, at the simplest level of description, over “surface” effects arising from interactions with the nuclear envelope. Thus, modelling the effects of interactions of LADS with the nuclear lamina by introducing weak monomer-specific interactions with the inner surface of the confining sphere in our simulations might be expected to modify the results we present here for specific chromosomes, but hopefully in a controllable manner. Second, we ignored nucleoli, formed around nucleolus organizer regions containing multiple copies of rRNA genes, with such regions located on the short arms of the acrocentric chromosomes 13,14,15, 21 and 22 (Németh and Längst, 2011). Third, we simulated the nucleus as a spherical shell containing our model chromosomes, although nuclear shapes exhibit considerable variability and much of the experimental data comes from experiments on the relatively flattened nuclei of fibroblasts (Bolzer et al., 2005). Our model could be generalised to account for the effects of variable nuclear shapes. Fourth, we ignored potential interactions *in trans* between chromosomes. Such interactions could potentially arise from the looping out of loci on different chromosomes to interact at transcription factories (Maharana et al., 2016). We could account for this by making designated monomers on different chromosomes “sticky” with respect to each other, thus coupling regions of different chromosomes that are known to physically localise together when co-transcribed. Fifth, in using RNA-seq data as a proxy for activity, we ignored non-coding transcription, since RNA-seq largely provides steady-state gene expression. Inferring activity from newer methods such as GRO-seq, which also extracts nascent and rapidly degraded transcripts, may help to provide a more accurate view of transcription-coupled activity. Last, the role of nuclear actin and associated motors remains unclear, although they could potentially contribute additional source of non-equilibrium noise (de Lanerolle, 2012). Indeed, all the possible improvements on our model that we list above could be incorporated, but only at the expense of more model detail and a number of further assumptions. These modifications of our model would have obscured the core argument of this paper, that the primary driver of many features of nuclear architecture is non-equilibrium activity that is inhomogeneous across chromosomes, so we choose to leave these questions to future work.

In first-principles approaches, a small set of initial model assumptions argued for on general grounds must yield consistent explanations and descriptions for all data, not just those the model abstracts in its construction. The advantage of simple models is that they enable us to concentrate on underlying principles that are often obscured by the complexity of real data, including intrinsic heterogeneities across cell populations, varied experimental and analysis procedures and the lack of sufficient statistics in some cases. Prior models for nuclear architecture in mammalian cells fail to reproduce many general attributes of nuclear architecture known from experiment. These properties – certainly their important trends – are emergent in our calculations, since they were not directly encoded in our model specification. This suggests that our methodologies provide hitherto unavailable biophysical insights into the determinants of large-scale nuclear architecture in metazoans.

## Methods

### Interactions of model chromosomes

Our model chromosomes (diploid, XX) occupy the interior of a spherical shell of radius *R*_0_. The interaction between neighbouring monomers is of the finitely-extensible non-linear elastic (FENE) form. These monomers further interact with (non-neighbouring) monomers via a Gaussian interaction, the Gaussian core potential used to model polymer brushes (Stillinger, 1976). Further details of these interactions and their bench-marking is discussed in Supplementary Information.

### Simulation Methodology

We adapt the widely-used LAMMPS code implementing Brownian dynamics (Plimpton et al., 2007) for a polydisperse polymer system with a local monomer-dependent effective temperature; for more details see Supplementary Information. For each monomer, LAMPPS applies a Langevin thermostat, via an over-damped equation of motion, with a different “effective” temperature *T_i_* associated to each monomer, reflecting its local level of activity. In thermal equilibrium, we have *T_i_* = *T_eq_* for all monomers.

### Units and Normalization

We work in de-dimensionalised units, discussed in Supplementary Information.

### Deriving effective temperatures

In the **gene density model**, the gene content of each 1Mb region is obtained from the GENCODE database (Harrow et al., 2012). Monomers containing a number of genes which fall below a preset cutoff are termed as “inactive” or “passive” and are characterized by an effective temperature *T* = *T_ph_* = 310K. Monomers possessing a larger number of genes are termed as “active” and assigned an effective temperature *T_a_* > *T_ph_*. For the **gene expression model**, we infer activities from transcriptome data, using FPKM values from processed RNA-seq output (Consortium et al., 2012). For the **combined model**, we use the same temperature assignments as for the **gene expression model** but, in addition, also take the top 5*%* of monomers by gene density as inferred from GENCODE, promoting them to a temperature of *T* = 12.

### Models for the looping of individual chromosomes

We use Hi-C data on GM12878, NHEK, IMR90, HUVEC and HMEC cells, obtained from data made publicly available by the authors of Ref. (Rao et al., 2014), to represent the effects of long-range looping within a chromosome. We ignore loops smaller than the 1 Mb scale since these are folded into our description of a single monomer. Across these cell types we have 236 (GM12878), 50 (NHEK), 116 (IMR90), 51 (HUVEC), and 13 (HMEC) loops that are larger than the 1Mb scale and which our model accounts for. These loops are represented by permanent FENE bonds, with an effective interaction strength that is the same as those of the springs which connect un-looped but connected monomers.

### Summary of Analysis

We calculate 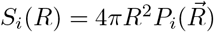, where 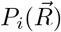 *dR* is proportional to the probability of finding a monomer of chromosome *i* at a radial vector 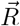 from the origin. For a uniform distribution, *S_i_*(*R*) = 4π*R*^2^. We compute *S_i_*(*R*) for every model chromosome indexed by *i*. We measure activity in successive radial shells by performing a configurational average over the effective temperature of every monomer in that shell. From these we extract a quantity similar to *S*(*R*), but normalize by 4π*R*^2^, so that the quantity plotted in the cut-away sphere representation simply represents the activity at radial distance *R*. The quantity S(R) measures the DNA density associated with a specific chromosome, across a radial shell at distance R from the nuclear centre, averaged over a large number of nuclei. The quantity S_*CM*_(*R*) measures a similar distribution, but of the chromosome centre of mass. We calculate the distribution of centres of mass of each individual chromosome similarly. To visually examine configurations we color-coded monomers belonging to individual chromosomes.

### Calculation of 3D shape of CTs

For each chromosome in our simulation, we draw a 3-d grid across the nucleus with a grid spacing of 0.2 – 0.6. We represent the monomers as spheres about which the density decays as a gaussian with defined width. Separating these monomers are cylindrical regions. The density about the axis of each cylinder is assume to fall off also as a Gaussian with specified width. The density at any given grid point, associated to a single chromosome, can then be computed by adding up the contributions from all spherical and cylindrical regions as defined above. Once such a density field is obtained, we can find the surfaces on which it attains a fixed value, the “implicit surface”. We adjust the scales governing the decay of the density distribution associated with the monomers and the cylindrical regions separating them as well as the associated constant specifying the implicit surface to optimise geometrical quantities associated with chromosome territories vis a vis experiments. Once fixed, these parameters remain the same for all chromosomes.

### Geometric properties of CTs

To calculate the two-dimensional properties of chromosome territories, we use the algorithm of Ref. (Sehgal et al., 2014). To compare our simulation data with data from 2d FISH, we project three-dimensional chromosome territories into the *xy* plane. We use the ellipticity calculations of (Žunić and Žunić, 2013). In the three-dimensional case, once we associate an implicit surface to a chromosome, that surface can further be triangulated using standard methods, such as the ISOSURFACE command in MATLAB. The total surface area of the chromosomes is obtained by adding the area of these triangles. To calculate the volume of the chromosome we count the number of grid points whose grid density values are more than the given isovalue density *c*. The asphericity Δ and shape (or prolateness) Σ parameters of a particular chromosome are calculated from the semi-axis lengths a, b, and c of the smallest ellipsoid that encloses all the monomers (Millett et al., 2009; Rawdon et al., 2008). The Khachiyan algorithm is used to find the smallest ellipsoid which encloses all the data points (Todd and Yildirim, 2007).

### Contact probability and contact map

The contact probability is computed using numerical calculations of the contact frequency of monomers of a given chromosome, averaged over a large number of configurational snapshots. When two monomers *i* and *j* of the same chromosome are separated in 3-d space by 2.5 units in terms of our scaled unit distance, we assume that they are in contact. If two monomers are in contact, they are close in distance. However, measures that look at the frequency of contacts will assign a larger frequency to such monomers which are predisposed to be close by.

## Author contributions

Conceptualization: GIM; Methodology: GIM, SS and AA; Formal Analysis and Validation: AA, NG and GIM; Investigation: AA and NG; Writing: Original Draft: GIM and AA; Writing - Review and Editing: GIM, AA, SS and NG; Visualization: AA and GIM; Software: AA and NG; Supervision: GIM; Project Administration: GIM

## Acknowledgments

We thank Ana Pombo, G.V. Shivashankar, Jean-Francois Joanny, Jacques Prost, Joe Howard, Sriram Ramaswamy, Mehran Kardar, B.J. Rao and Frank Jülicher for useful discussions. We are especially grateful to Kundan Sengupta, Sandhya Koushika and P. Ranjith for detailed input and discussions regarding our manuscript. The work of GIM was supported in its early stages through a DAE-SRC Outstanding Researcher Fellowship. We acknowledge the use of the Annapurna HPC at IMSc.

## Supplementary Information

### S1 Model Overview

We model individual chromosomes as polymer chains, composed of spherical monomers representing consecutive 1Mb sections of each chromosome. There are 23 pairs of chromosomes, describing a human diploid female (XX) cell. These model chromosomes are confined within a spherical shell representing the nucleus. The monomers are connected through nonlinear springs and interact via a finitely-extensible nonlinear elastic (FENE) interaction (Kremer and Grest, 1990). Individual monomers are self-repelling and the total number of monomers is 6086 for a diploid XX human cell. Monomers experience forces from other monomers, arising from both bonded and non-bonded interactions. Additionally, each monomer experiences random forces arising from thermal as well as active fluctuations. We treat such active noise as analogous to thermal noise, drawing particular realisations of the noise from a Gaussian distribution with zero mean and a variance set by the effective temperature.

To represent activity we assign an active temperature to monomers in three ways. In the simplest model, the “gene density” model, the temperature assigned to each monomer reflects the gene density associated with the specific region of the chromosome associated to that monomer. The gene content of each such 1Mb region is obtained from the GENCODE database (Harrow et al., 2012). We label each monomer as active or inactive, associating one of two temperatures (*T* = 1, 12) to the monomer depending on its label. Temperatures are scaled in units of physiological temperature *T_ph_* = 310*K*.

A second model, the “gene expression” model, assumes that the temperature assigned to each monomer is proportional to the amount of RNA transcript generated across that region of chromosome. Transcript levels then provide a proxy for the intensity of active processes locally. We infer these levels using transcriptome data, as described below. Both the gene expression and gene density models fit specific aspects of the experimental data well. However, a model description which appears to provide the most comprehensive fits to the data combines features of both these models. The model used in this paper, the “combined model”, bases itself largely on the gene expression model but also assigns high activity to a fraction of monomers with the highest values of gene density.

An important part of our model is the incorporation of existing prior information regarding the looping of chromosomes. Here, we use data from Hi-C experiments to assign permanent loops to our model chromosomes. We simulate our system using standard Brownian dynamics methods. After verifying that the system has achieved a non-equilibrium steady state, we compute all properties of interest, including the distribution of DNA density and of chromosome centre of mass, territorial organisation, shape statistics and spatial distance maps from which we can infer potential contacts.

#### S1.1 Interactions of model chromosomes

Our model chromosomes occupy the interior of a spherical shell. The radius of this shell is *R*_0_, which we take to be 17.2 in the reduced units we derive below. The interaction between neighbouring monomers (labeled as *i*, *i* + 1, with position coordinates **r**_*i*_, **r**_*i*+1_) is of the FENE form

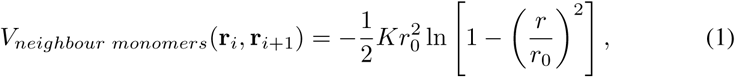

where *K* is a spring constant and *r*_0_ is the maximum length of the bond; in contrast, our earlier work used the simpler harmonic spring interaction (Ganai et al., 2014). These monomers further interact with (non-neighbouring) monomers via a Gaussian interaction, the Gaussian core potential used earlier to model polymer brushes (Stillinger, 1976)

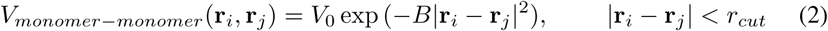

The effective pair potential at zero separation, *V*_0_, is chosen to be of order *k_B_T_ph_*, with *k_B_* the Boltzmann constant and *T_ph_* the physiological temperature: *B* = 1.0, *V*_0_ = 1.5 and *r_cut_* = 3.5. The interaction between each monomer and the confining sphere vanishes if the monomer centre falls within the sphere. Outside the sphere and within a cutoff *r_c_*, the monomer experiences a Lennard-Jones potential that diverges as the distance to the cutoff is reduced. The parameters are ∊ = 250, σ =1 and *r_c_* = 1 for this Lennard-Jones potential. We take *r_0_* = 10 and *K* = 4.17, for both bonded neighbors and non-bonded monomers connected by long range loops.

#### S1.2 Simulation Methodology

Our numerical evolution of the system of monomers adapts the widely-used LAMMPS code implementing Brownian dynamics (Plimpton et al., 2007) for a polydisperse polymer system with a FENE interaction between monomers and a monomer-specific effective temperature. The effective temperature is incorporated as a local monomer-dependent effective temperature. For each monomer, LAMPPS applies a Langevin thermostat, via the following over-damped equation of motion,

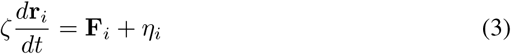

where **r**_*i*_ represents the location of the *i^th^* monomer, ζ is a drag coefficient, **F**_*i*_ accounts for all non-stochastic forces acting on the monomer and *η*_*i*_ represents stochastic forces (gaussian, with vanishing cross-correlations) arising from both active and thermal fluctuations. The noise is assumed Gaussian distributed, with cross-correlations vanishing at all times irrespective of monomer labels. The diagonal correlators, at equal times and for the same monomer, are non-zero and obtained from 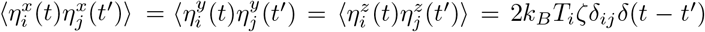. Here *T_i_* is an “effective” temperature associated to each monomer, reflecting its local level of activity. We represent each of the components of 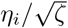 as the product of a Gaussian random number with zero mean and unit variance with the quantity 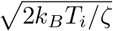. In thermal equilibrium, we have *T_i_* = *T_eq_* for all monomers. The largest number of monomers (249) are found in Chromosome 1 and the smallest (47) in Chromosome 21.

In comparison to the model in Ref. (Ganai et al., 2014), the model described here has 12 fewer monomers, a consequence of differences between the GeneCards and GENCODE databases.

#### S1.3 Units and Normalization

Following Refs. (Kreth et al., 2004; Ganai et al., 2014), we consider a chromatin volume fraction 0.1 ≤ *ϕ* ≤ 0.2. The monomer is assumed to have a diameter *d* ⋍ 500*nm*; the equilibrium domain separation is *l*_0_⋍ 600*nm*. Both these quantities accord with computed Kuhn lengths of ≈ 300*nm* (Kreth et al., 2004; Rosa and Everaers, 2008). Assuming that the radius of the nucleus is *R_0_* ⋍ 8.6*μm* yields a packing fraction of *ϕ* ⋍ 0.15. We ignore the marginal differences in nuclear volume across cell types; such volumes differ by at most a factor of 1.5 for the cell types we consider. We scale all lengths in units of *d* and measure energies in units of *k_B_T_eq_*. All chromosomes are fairly tightly confined to *R*_0_. We can choose units of time (τ) such that ζ = 1. We can approximate the value of ζ appropriate for this calculation from the Stokes relation: ζ ⋍ 6*πη_S_R* where *R* is the hydrodynamic radius appropriate to the monomer size. Assuming that the appropriate value of the viscosity at such scales is *η_s_* ~ 10*η_w_* with *η_w_* the viscosity of water (8.9=10^−4^*P*_*a.s*_), its numerical value is then ζ = 8.38=10^−8^*N .s/m*. With this choice, τ is then (500.0)^2^ζ/6*k_B_T* ‘ 8.16 = 10^−1^*s*. Since τ ≈ 10^−1^*s*; our choice of time-step of 0.001 thus corresponds to real-time evolution by 10^−4^s.

#### S1.4 Deriving effective temperatures

##### 51.4.1 Gene density model

In the gene density model, the gene content of each 1Mb region is obtained from the GENCODE database (Harrow et al., 2012). Monomers containing a number of genes which fall below a preset cutoff are termed as “inactive” or “passive” and are characterized by an effective temperature *T* = *T_ph_* = 310*K*. Monomers possessing a larger number of genes are termed as “active” and assigned an effective temperature *T_a_* > *T_ph_*, argued for thus: The hydrolysis of a single ATP molecule yields approximately 20*k_B_T_ph_* of energy, some part of which is transduced into local mechanical work with the remaining part begin dissipated. Thus, the characteristic energy scales associated with such hydrolysis events, modelled in terms of an effective temperature, must be some reasonable fraction of this scale. We have experimented with several different choices of *T_a_* as well as the cutoff, finding that a relatively small spread between physiological and active temperatures is sufficient to generate the activity-dependent biophysical structuring this paper discusses. For concreteness, here we take the maximum value for the active temperature to be *T_a_* = 12, where *T_a_* is measured in units of *T_ph_*. (Measurements of the diffusion constants of individual gene loci in bacteria and yeast provide evidence for a similar spread in local “effective” temperatures, as inferred from an Einstein relation, and that a related variation in local active forces has been suggested to explain observations from colloidal micro-rheology in the nucleus (Zidovska et al., 2013; Weber et al., 2012; Hameed et al., 2012))

##### 51.4.2 Gene expression model

For the gene expression model, we infer activities from transcriptome data. We downloaded processed RNA-seq data from the ENCODE project website (Consortium et al., 2012). The data provides FPKM (Fragments Per Kilobase of transcript per Million mapped read, with fragment referring to a pair of reads for paired-end data) values that quantify transcripts generated across the human genome. We used five cell types GM12878, HMEC, HUVEC, IMR90 and NHEK for our analysis. We consider all genes whose FPKM value lies above a specified cutoff to reduce noise in the data for each cell type. We then summed the FPKM value for all these genes whose chromosome position (mid position of start and end coordinate of a gene) lies within our 1 Mb interval to assign an activity value to that monomer. We went on to assign effective temperatures proportional to such activity values using a derivative cutoff method. We examined the expression data discussed above, sorted it in increasing order of expression, and then took a numerical derivative of this data, using the *diff* function of MATLAB. We set a cutoff of 0.2 for this slope but our results were largely insensitive to where this cutoff was chosen. Given the generic shape of the plot for the sorted activity, which has two regions in which activity increases sharply separated by a plateau region, this procedure automatically yields two cutoff lines and three demarcated regions. Monomers in the plateau region are assigned a constant value of active temperature, between *T_ph_* and the maximum value of the active temperature 12*T_ph_*.

For each cell type, given similar curves of sorted expression value, such cutoffs on the data represent the effects of activity on each monomer. We can translate this into an active temperature. In (Figure 1H), for each cell type, we show these cutoff lines. We assign the lowest temperatures to monomers whose activity falls below the value it takes in the plateau region. Monomers associated with the plateau are assigned a common temperature of 6. Finally, monomers with the highest expression values and thus the highest activity, are assigned a temperature which interpolates, in units of 1, between 7 and 12. The effects of variation of activity are strongest for these monomers, as is reasonable since activity increases steeply in this region.

##### S1.4.3 Combined model

For the combined model, we use the temperature assignments indicated above. However, in addition, we also take the top 5% of monomers by gene density as inferred from GENCODE, promoting them to a temperature of *T* = 12.

#### 51.5 Summary of Analysis

Our simulations are run for 10^7^ time steps, with around 4 × 10^6^ steps discarded to ensure adequate equilibration. All data are averaged over at least 25 independently initialised configurations, with each initial configuration contributing 6000 independent measurements as the simulation proceeds. We verified that the same steady state properties were achieved irrespective of initial (random) configuration. Since the probability of finding a chromosome at a radial separation r from the origin depends only on the modulus of r, *i.e*. |r| ≡ *r*, we calculate the probability of finding a monomer belonging to a specific chromosome at a radial distance from the origin, for each chromosome. The main text describes the different distribution functions we calculate.

#### 51.6 Calculation of 3D shape of CTs

We begin by drawing a 3-d grid across the nucleus with a coarse grid spacing of 0.5 in our scaled units. We calculate the DNA density associated with our polymer model for chromosomes on this grid as described below. Chromosome densities are then defined at the grid points for each simulation configuration. We represent chromosome configurations in the following ways. First, individual monomers are considered as spheres about which the density decays as a normalized gaussian in the radial variable. The characteristic scale of the Gaussian is set by the length scale *R_s_*. We experimented with various choices of *R_s_* to see what might best represent the data. Between these monomers, we assume that the DNA configuration can be described as a cylinder with fixed radius, with the DNA density about the axis of each cylinder assumed to fall off also as a Gaussian with specified width *R_c_*. We have varied *R_c_* to obtain optimal fits as well, but finally fixed on *R_c_* = *R_s_* = 2σ as an optimal choice for all chromosomes across all configurations and all cell types.

The density at any given grid point can then be computed by adding up the contributions from all spherical and cylindrical regions associated to a chromosome. We then interpolate these density values to a smaller grid, typically of spacing 0.17 using MATLAB’s INTERP3 function. Once such a density field is obtained, we can find the surfaces on which it attains a fixed value, the “implicit surface”.

Our calculation of the density at a given grid point proceeds as follows. Consider the location of the grid point *O_j_*, where *j* indexes the specific grid point. Let *n* be the length of chromosome *C^k^* and denote the *i*th monomer of the k^*th*^ chromosome as 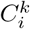. The visual computational of such calculation is shown in (Figure S16). The two consecutive monomer 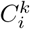 and 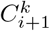 of *k* chromosome is represented by small circles in, and lines representing the centreline of the associated cylindrical regions. *O_j_* is the grid point where we have to calculate the spherical density *D_sphere_* which is contributed by mentioned two monomers shown in (Figure S16A) and similarly contribution by their centerline in form of cylindrical density *D_cylinder_* is shown in (Figure S16B). The density at the grid point *O_j_* arising from chromosome *k* is then computed from the following formula and also shown in (Figure S16C).

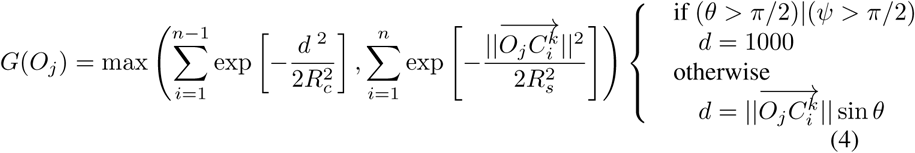

Here 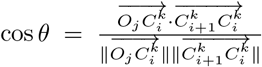 and 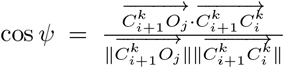. These condition sensure that (a) the contribution of the associated cylinder to the density at any grid point is cylindrically symmetric about the line joining neighbouring monomers and that (b) the contributions from the spherical regions is also accounted for.

After the density values at all grid points is computed, the isosurface command from MATLAB is used to draw the implicit surface (*F*(*x*, *y*, *z*) = *c*) for the chromosome given a density isovalue *c*. A smaller value of *c* yields a loose cloud-like surface around chromosomes while larger values of *c* gives tighter, more well-defined surfaces around them. We chose a value of *c* such that the chromosome territories it yields are visually equivalent to those obtained in experiments based on similar iso-surface representations of experimental FISH data. The 3d surface representation of such chromosome is shown in (Figure 1E). This computation is used in the fits to the experimental data shown in (Figure 5C).

#### S1.7 Calculation of 2D shape of CTs

To calculate the two-dimensional properties associated to projected chromosome territories. To compare our simulation data with data from 2d FISH as obtained in those experiments and other similar ones, we project three-dimensional chromosome territories into the *xy* plane for specificity, noting that averaging over configurations which are rotationally symmetric implies that all projections should be equivalent. A schematic illustrating this is shown in (Figure 1G). The calculations of Ref. (Sehgal et al., 2014; žunić and žunić, 2013) rely on the computation of the geometric moments *M* of the image *I*. For calculating the ellipticity and regularity of 2d CTs image, we first normalize, scale and rotate the complementary binary image *I* of our projected chromosome territories such that

1. The area of shape is 1
2. The centroid of shape coincides with the origin and
3. The orientation of shape is 0 (implying that the long axis is parallel to the x-axis).

The scaled moments then are:

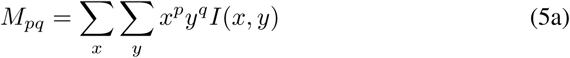

Where I(x,y) is the pixel intensitites of the grayscale image and {*p, q*} can each be drawn from 0 … ∞, and refer to the order of moments. We choose each of {*p, q*} from 0,1,2 for specificity.

Given the moments, we compute the ellipticity ∊ from

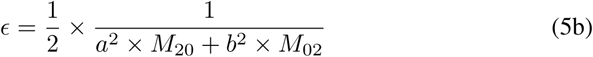

where *a* and *b* are given by

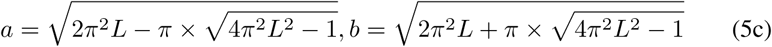

and *L* = *M*_20_ + *M*_02_

As ∊ tends to 1, projected chromosome territories resemble ellipses. To calculate the regularity of these projected territories, the area ratio of each CT over its convex hull is determined, using the MATLAB functions BWAREA and BWCONVHULL. If a chromosome is regular, this ratio should be close to 1. As the irregularity in projected CTs increases, this ratio will decrease.

#### S1.8 Methods for calculating 3d shape statistics for chromosome territories

We use two methods for finding the shape statistics of chromosomes. In the ellipsoidal fit method, the center, rotation, and principal radii of the smallest ellipsoid which encloses all the data points of a polymer are calculated using Khachiyan algorithm (Todd and Yildirim, 2007). An alternative grid-based method is described in the subsection **Calculation of 2D shape of CTs**. The ellipsoid fit method is an easy and fast method for finding the approximate shapes of the polymer, but is less accurate than the grid-based method. On the other hand, the grid-based method is computationally expensive. In our work, we calculate the shapes of 2d and 3d territories using the grid-based method but use the simpler ellipsoidal fit method for the shape parameters Δ and Σ described below.

The difference in the computation of chromosome territory properties between the grid-based method and the ellipsoid fit method is illustrated in (Figure S17).

#### S1.9 Asphericity Δ and shape Σ parameter calculation of chromosomes

The asphericity Δ and the shape parameter Σ of individual chromosomes can be calculated from the 3 semi-principal radii (a,b,c) of the ellipsoid obtained for each ellipsoidal fit to a chromosome territory. The asphericity Δ parameter given in Eq. 6a characterizes the average deviation of the chain conformation from spherical symmetry (Millett et al., 2009; Rawdon et al., 2008). The shape Σ parameter measures the prolateness or oblateness of chromosomes and is defined in Eq. 6b.

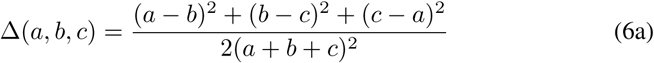

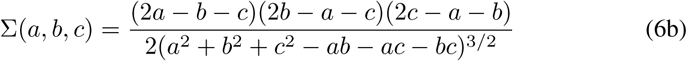

The parameter Δ is bounded in the regime 0 ≤ Δ ≤ 1. The Δ value is 0 for the perfect sphere when (*a* = *b* = *c*) and 1 for rod shape when (*b* = *c* = 0). The shape or prolateness parameter Σ is bounded by −1 ≤ Σ ≤ 1. The Σ is −1 for perfect oblate shapes, e.g. when (*a* = *b* > *c*) and 1 for perfectly prolate shapes, e.g. when (*a* > *b* = *c*), providing a useful index for chromosome shapes.

---

**Figure S1:**
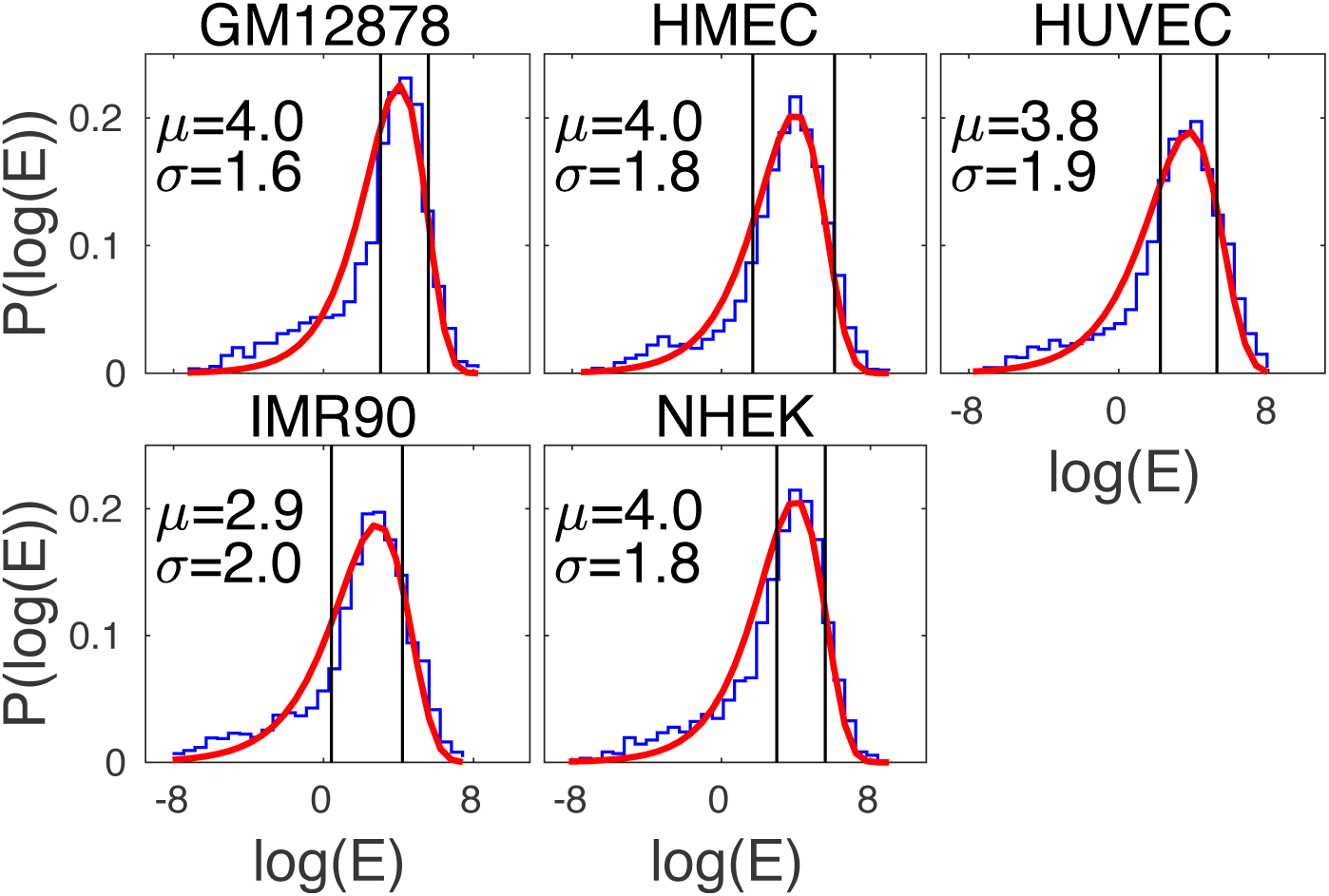
Distribution of expression values derived from transcriptome data. The histogram of the log of gene expression values as obtained from transcriptome data for 5 cell types is shown in blue. Each cell type name is provided at the top of each subfigure. The sub-figures illustrate a fit of histogram values to an extreme value distribution, the Gumbel distribution, shown in red. The Gumbel distribution for a single variable *x* takes the form 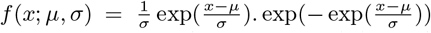. The fitting parameters μ and σ for the Gumbel distribution in each case are provided in each subfigure. Two black vertical lines, derived from the analysis that led to (Figure1H), demarcate the histogram into 3 regions. The distribution has a long tail towards low expression values and a more sharply decaying right tail towards high expression values.

**Figure S2:**
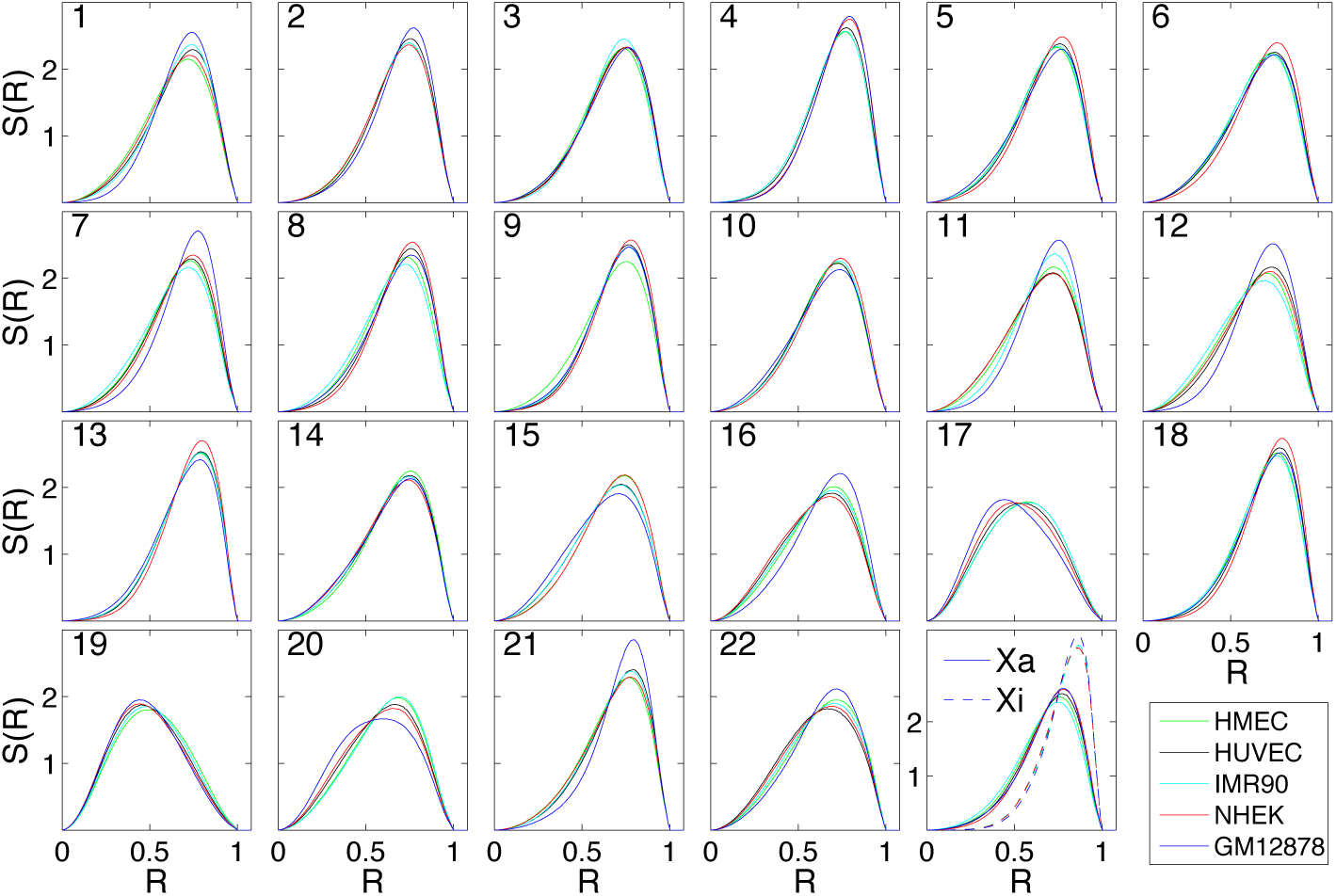
Computed Distribution functions S(R) for all simulated chromosomes across 5 cell types. Distribution functions S(R) for each simulated chromosomes for GM12878 (blue), HMEC(green), HUVEC(black), IMR90(cyan) and NHEK(red) is shown. The active chromosome Xa and inactive chromosome Xi are shown with solid and dashed lines. Chromosome numbers are mentioned in the left upper corner of each subfigure.

**Figure S3:**
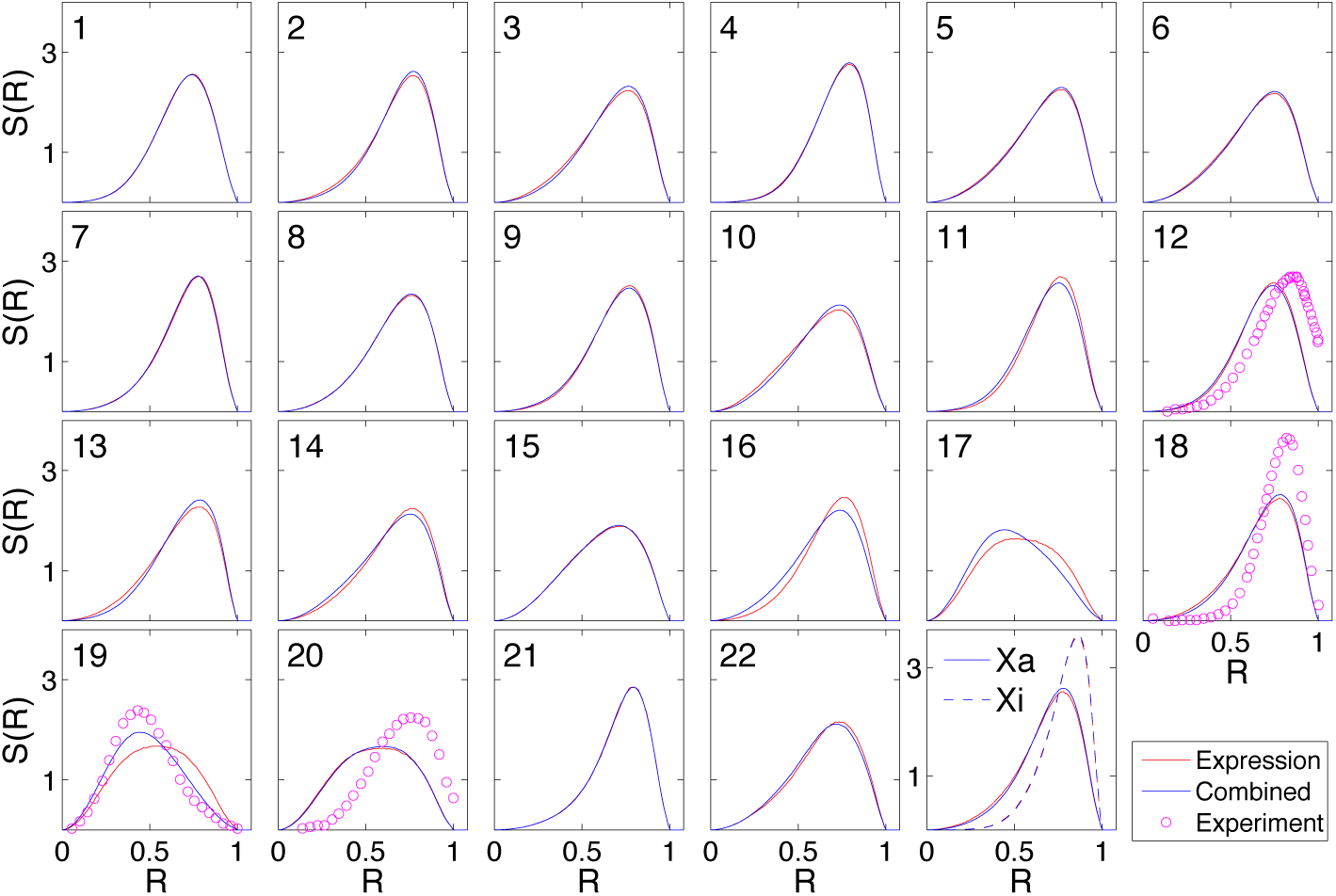
Distribution functions S(R) for all chromosomes, within the “gene expression” and “combined” models. S(R) for each chromosome within the gene expression (red) and the combined model (blue) are shown. Experimental data from Ref. (Kreth et al., 2004) for Chromosomes 12, 18, 19 and 20 are shown as magenta-coloured circles.

**Figure S4:**
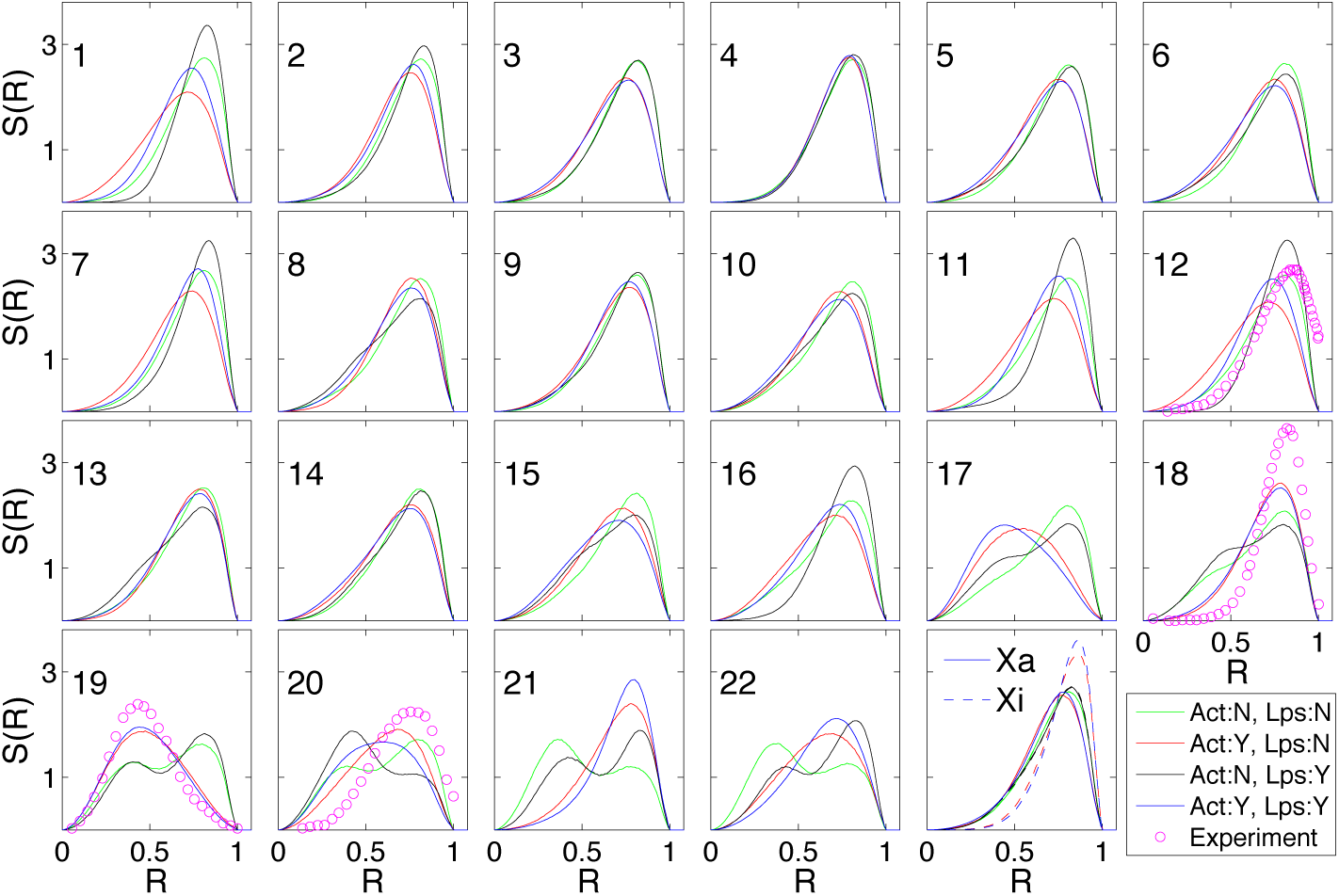
Distribution functions S(R) for all chromosomes with different combinations of activity and loops. The monomer density distribution for each chromosome is shown for the models mentioned below. All cases involving activity are shown for the combined model. **Act:N, Lps:N** Both activity and loops are switched off, with all monomers at the same effective temperature of T = 1, shown in green; **Act:Y, Lps:N** Activity is present, implying an inhomogeneous distribution of temperatures, but loops are switched off, shown in red; **Act:N, Lps:Y** Activity is absent but loops are present, shown in black; **Act:Y, Lps:Y** Both activity and loops are present, shown in blue color. This is the original “combined model”, also shown in (Figures 3A and 3B) and (Figure S3); The experimental data for chromosomes 12, 18, 19 and 20 are shown as magenta-colored circles, using data from Ref. (Kreth et al., 2004).

**Figure S5:**
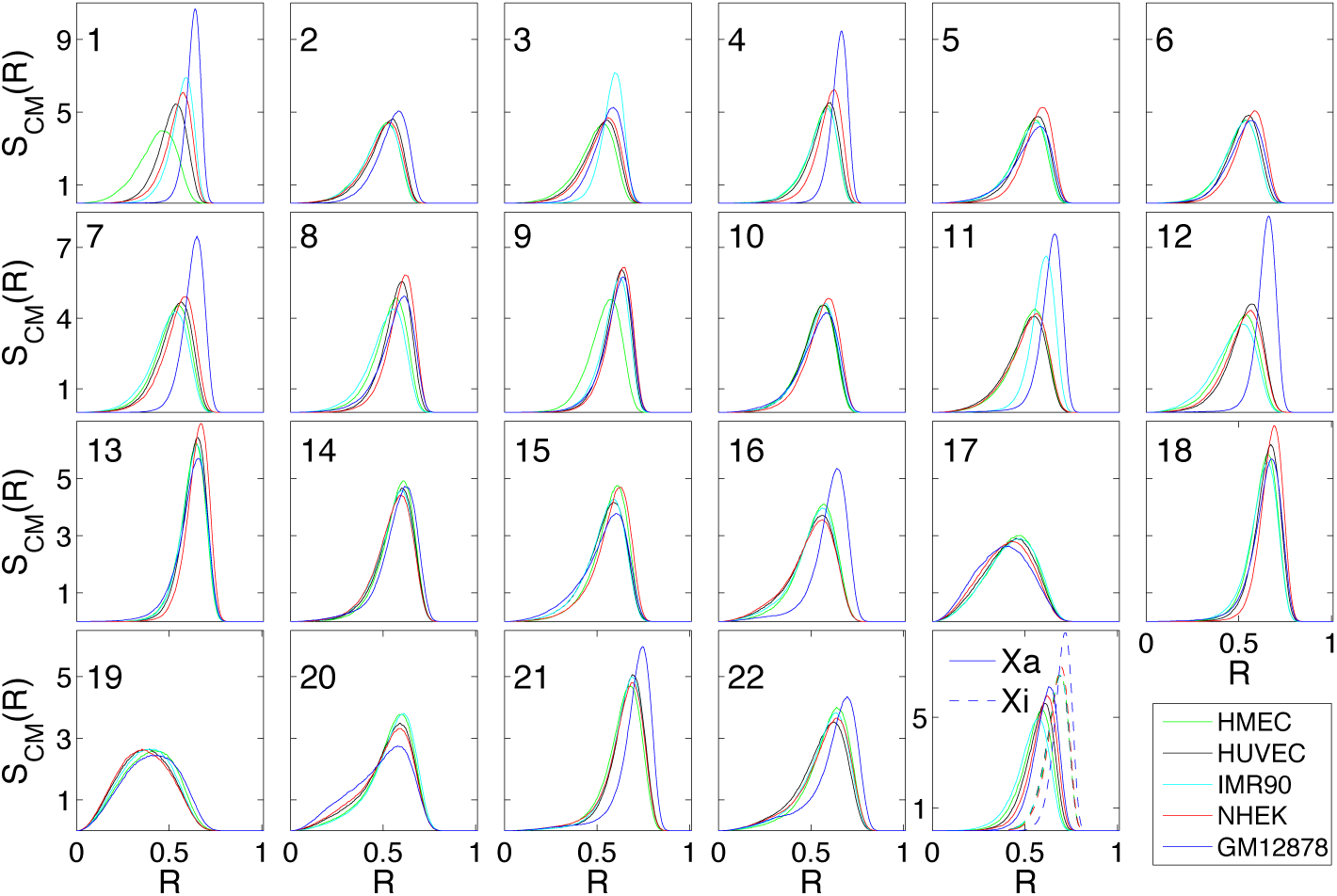
Centre of mass distributions S_*CM*_(*R*) for all simulated chromosome in 5 cell types. The centre of mass distribution S_*CM*_(*R*) for each simulated chromosome for GM12878(blue), HMEC(green), HUVEC(black), IMR90(cyan) and NHEK(red) is shown.

**Figure S6:**
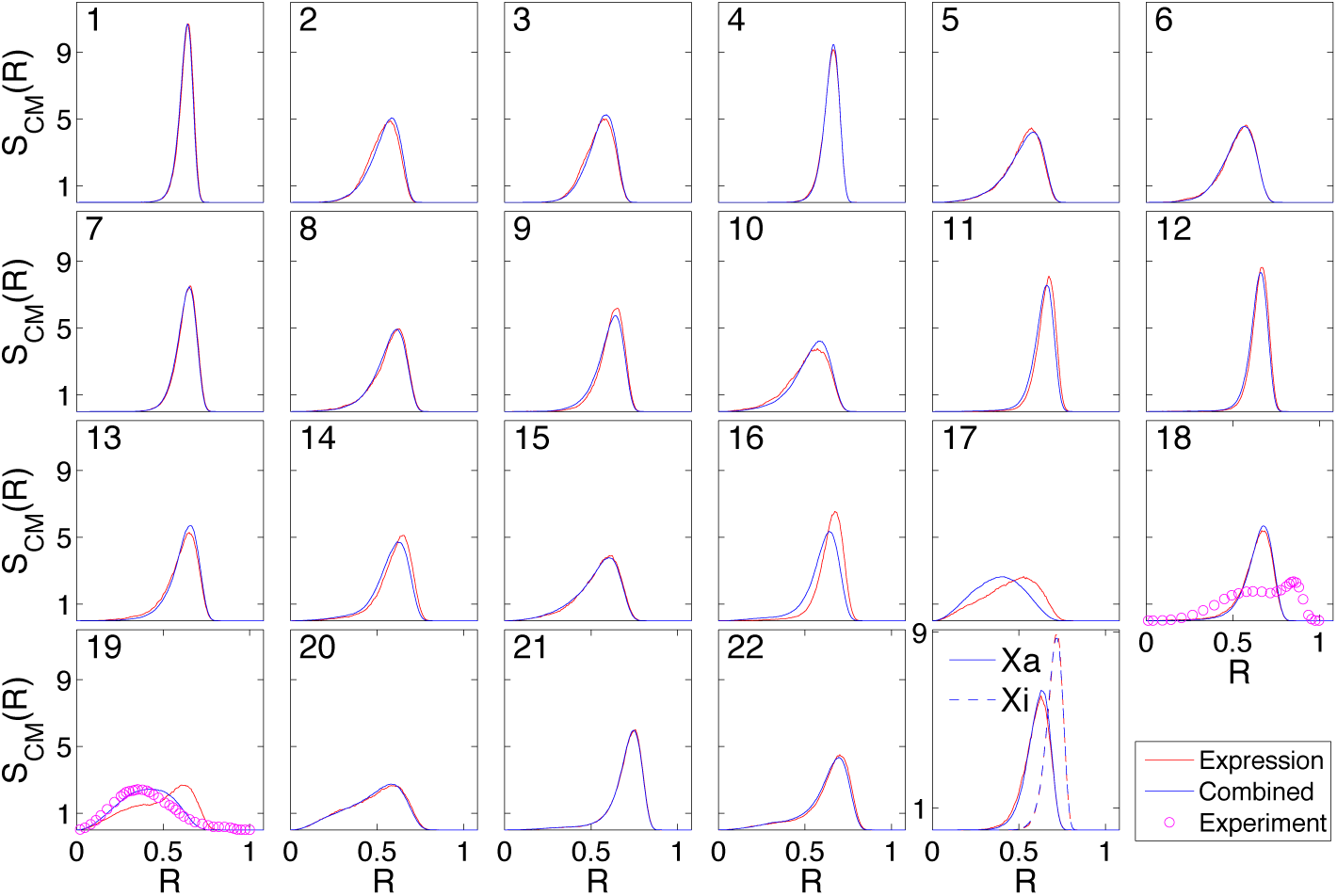
The centre of mass distribution S_*CM*_(R) for all chromosomes, within the “gene expression” and “combined” models. S_*CM*_(R), the centre of mass distribution of each chromosome for gene expression (red) and combined model (blue). Experimental data from Ref. (Kalhor et al., 2011) for Chromosomes 18 and 19 are shown in magenta.

**Figure S7:**
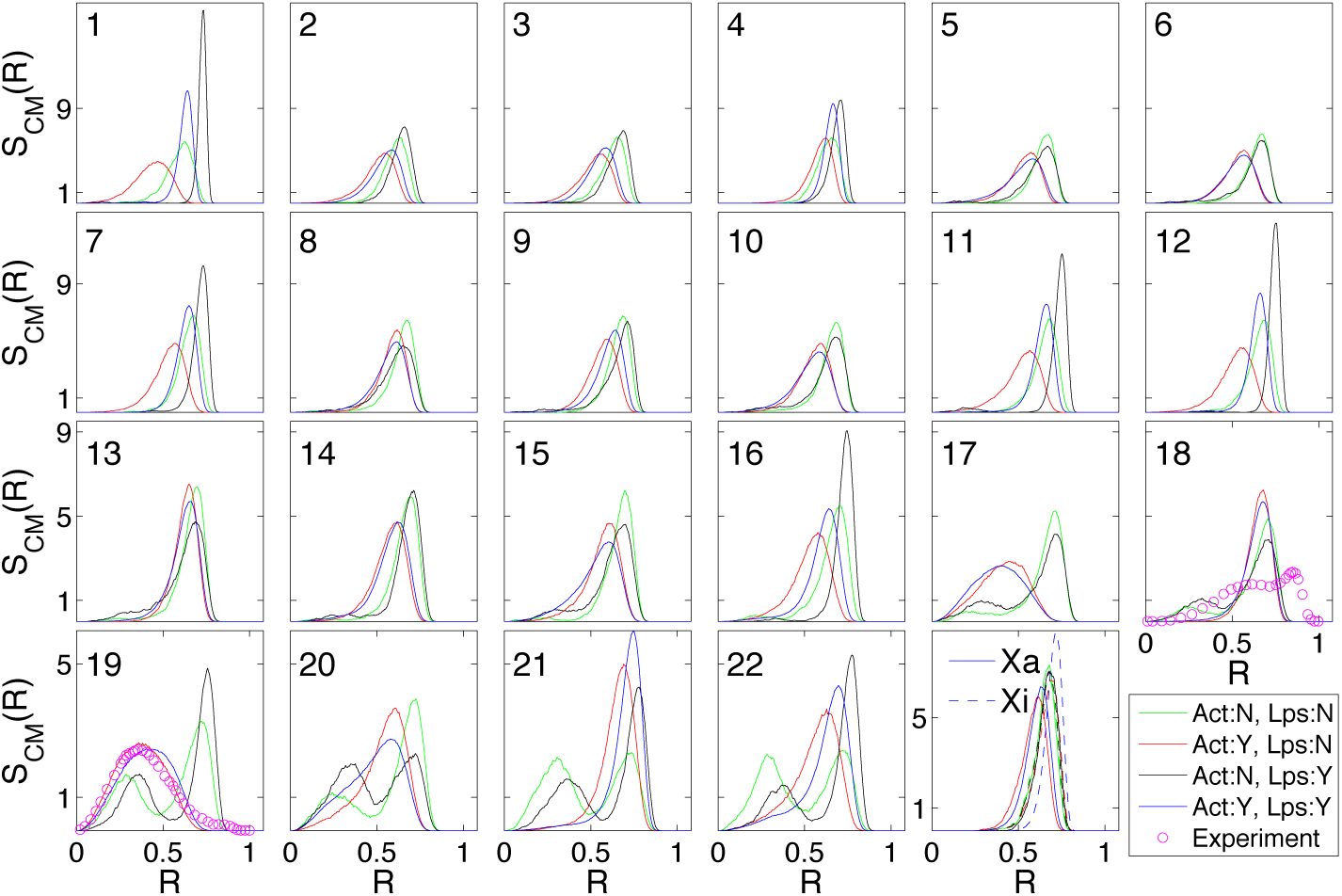
Centre of mass distribution S_*CM*_(R) for all chromosomes with different combinations of activity and loops. S_*CM*_(R) center-of-mass distribution for each chromosome is shown for the models described below. **Act:N, Lps:N** Both activity and loops are switched off, with all monomers at the same effective temperature of T = 1, shown in green; **Act:Y, Lps:N** Activity is present but loops are switched off, shown in red; **Act:N, Lps:Y** Activity is absent but loops are present, shown in black; **Act:Y, Lps:Y** Both activity and loops are present, shown in blue color. This is the original “combined model”, also shown in (Figures 3C and S6); The experimental data for chromosomes 18 and 19 are shown using magenta circles using data from Ref. (Kalhor et al., 2011).

**Figure S8:**
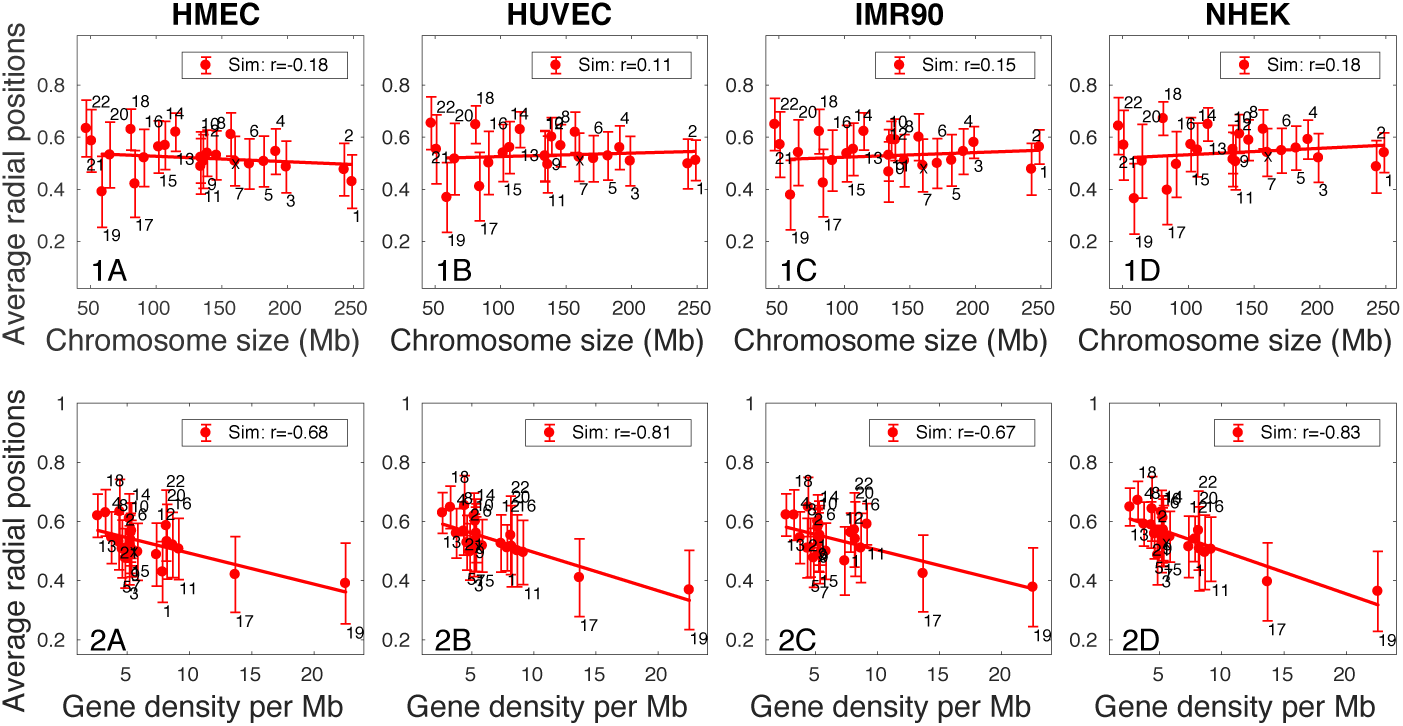
Mean centre-of-mass location for each chromosome against chromosome sizes and chromosome gene density. Simulation prediction for the mean center of mass location of HMEC, HUVEC, IMR90 and NHEK cell types, for each chromosome as a function of chromosome size in the upper row and chromosome gene density per Mb in the lower row. For the fit against chromosome size, Chr 21 and 22 are excluded. The relative radial position 0 and 1 represent the center and periphery of nucleus respectively. Chromosome numbers are indicated above or below each error bar. The Pearson correlation coefficient (PCC) *r* between chromosome size/gene density & average radial positions are mentioned in each subfigure Simulation points are fitted to a straight line accounting for all chromosomes. As can be see, the correlation to chromosome size is weaker than the correlation to chromosome gene density.

**Figure S9:**
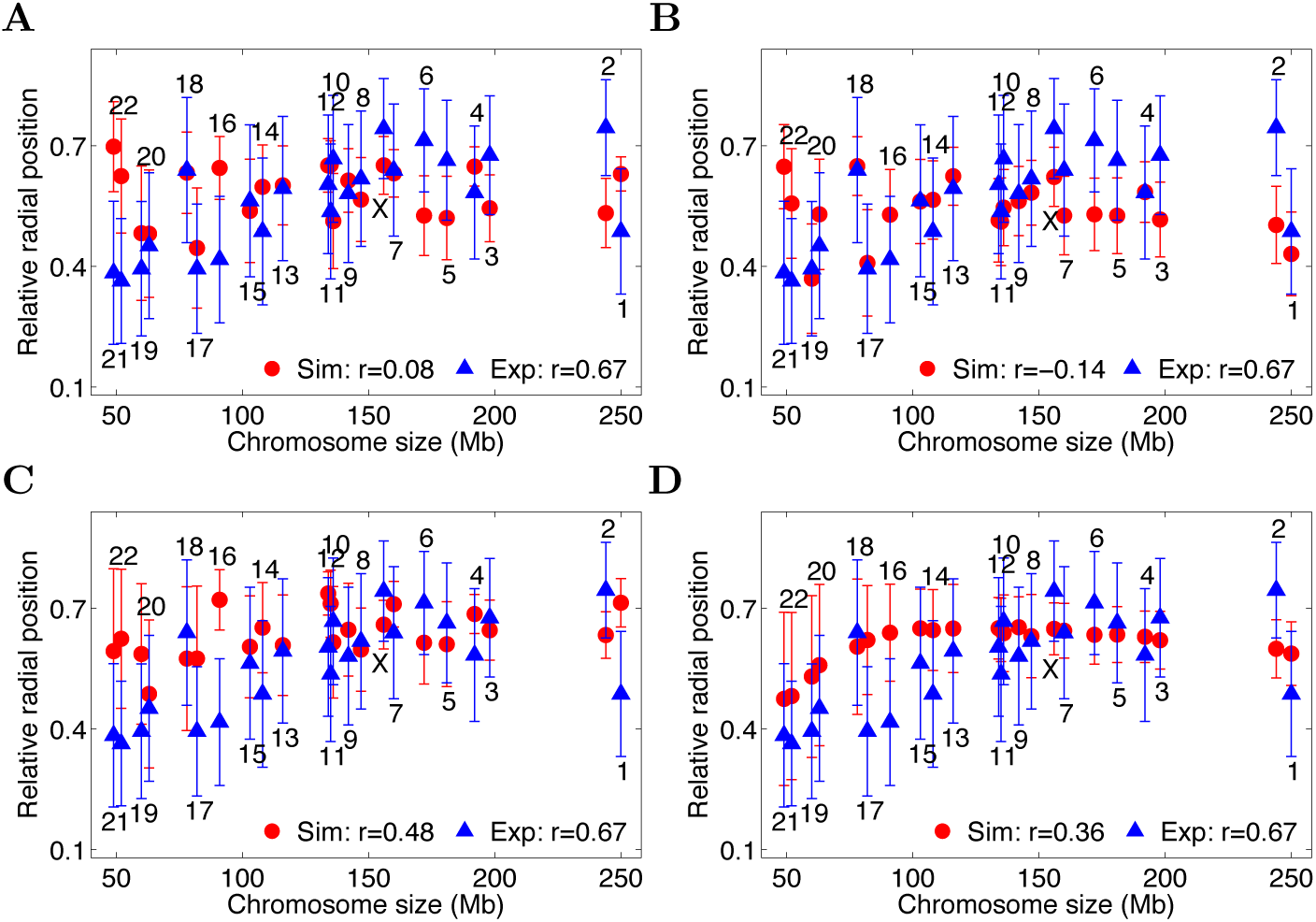
The relative centre-of-mass position of each chromosome for different combinations of activity and loops. The relative centre-of-mass position of each chromosome, in increasing order of size, shown for different models; **(A) Gene expression** The effective temperature assignment of monomers is derived from gene expression model of GM12878 cell type and loops are extracted from Hi-C data; **(B) Act:Y, Lps:N** The effective temperature assignment of monomers is taken from the combined model of GM12878 cell type but no loops are present; **(C) Act:N, Lps:Y** All monomers are at the same temperature (no activity), but loops inferred from Hi-C are present; **(D) Act:N, Lps:N** All monomers are at the same temperature (no activity) and loops are also absent; Simulation data points (red oval) are shown together with the experimental data (blue triangle) (Kalhor et al., 2011) along with respective error-bars. Chromosome numbers are mentioned above or below each error-bar. Pearson correlation coefficients *r* illustrate the quality of linear fits to the data and are shown for both simulation data and experiments. Experimental data are extracted from Ref. (Kalhor et al., 2011).

**Figure S10:**
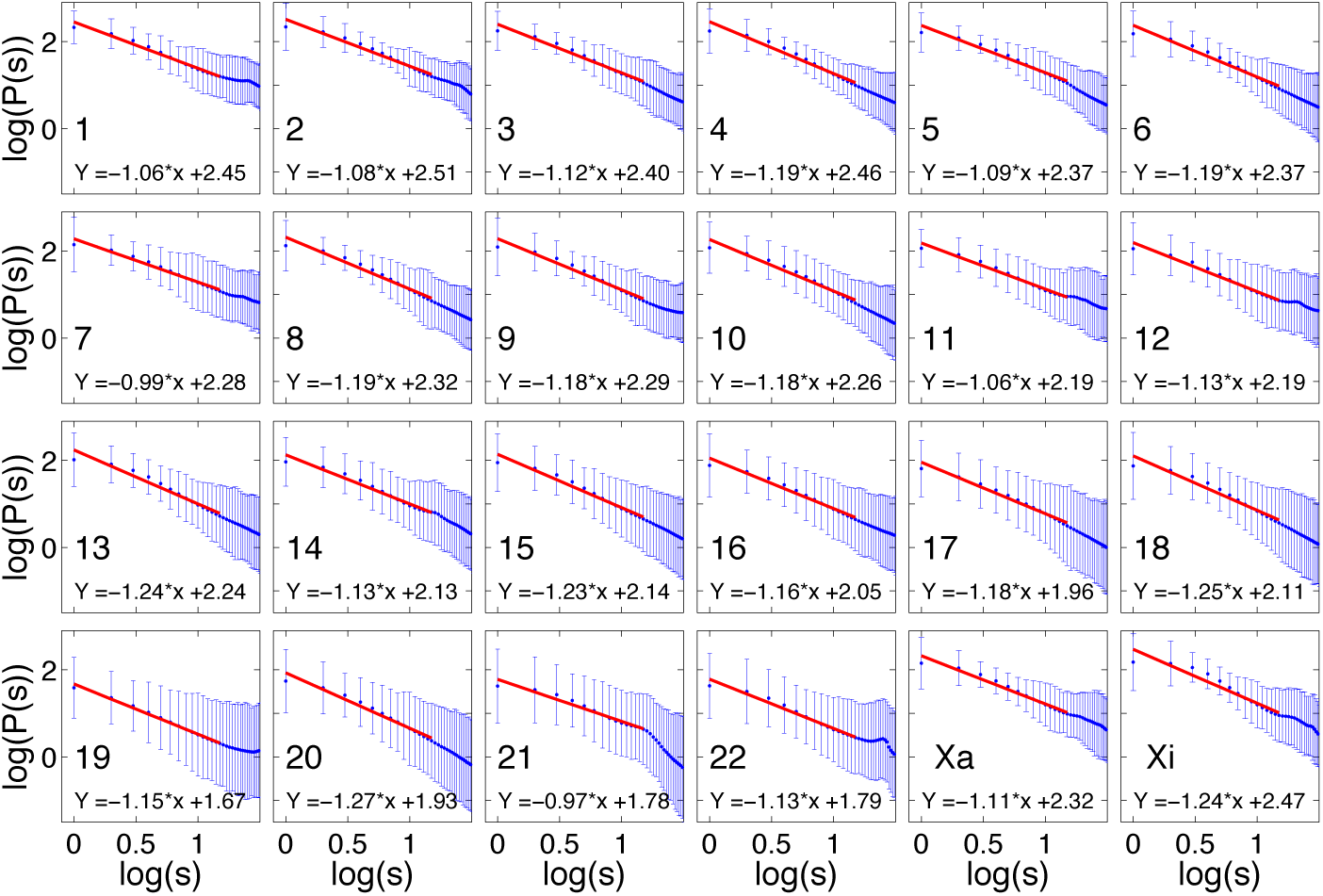
Contact probability *P*(*s*) for each chromosome in the GM12878 cell type. The scaling of monomer contacts within the initial 10 Mb region for each chromosome is shown with blue dots, accompanied by error-bars. The best power law fit to the data is provided along with each subfigure, in red. The chromosome number is mentioned in the bottom corner of each subfigure. The range of exponents provided by the fit is 0.97 – 1.27. The plot shown here exclude superloops for the Xi. (These are discussed in the main text and figures.)

**Figure S11:**
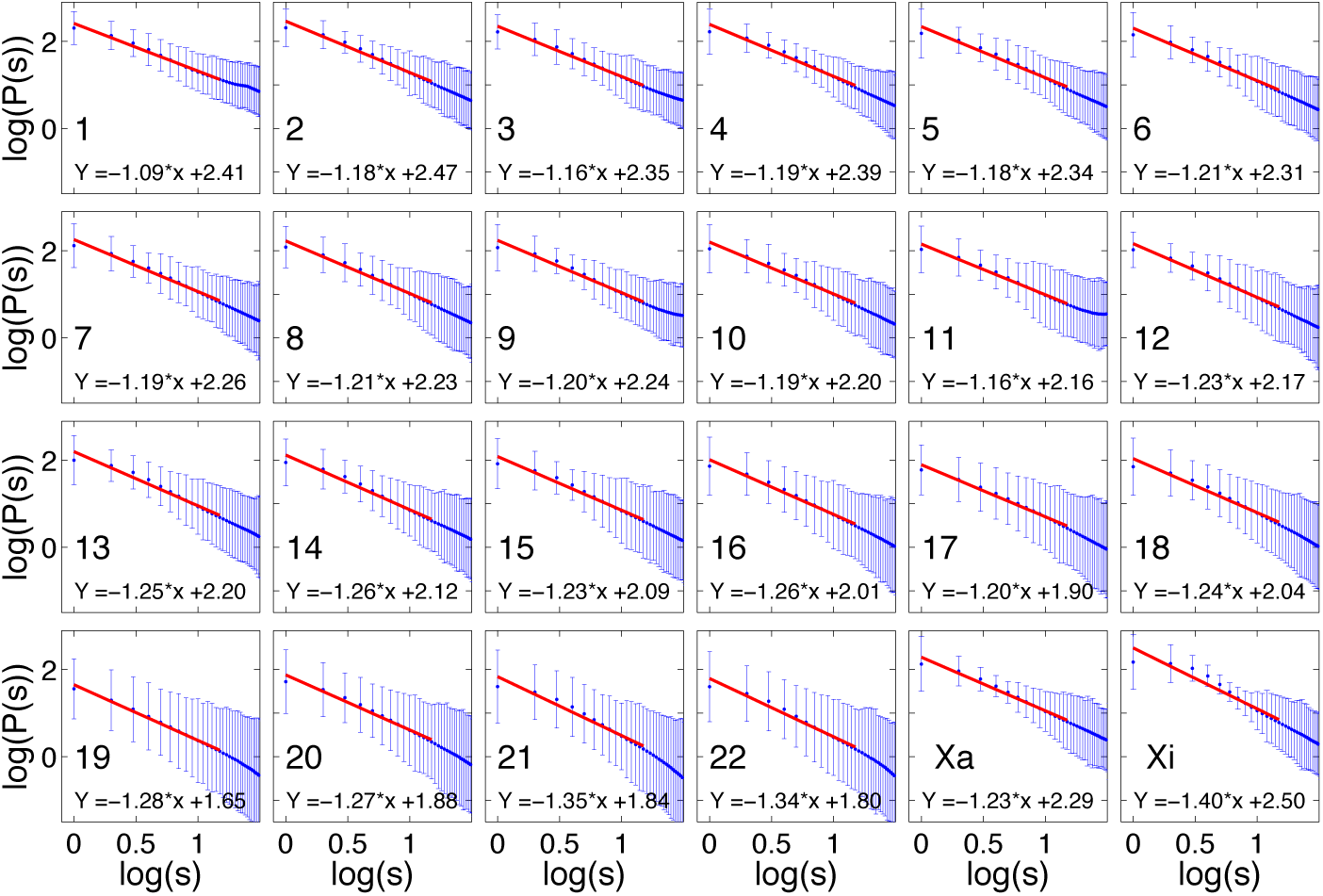
Contact probability *P*(*s*) for each chromosome in the IMR90 cell type. The scaling of monomer contacts in the initial 10 Mb region for each chromosome is shown with blue dots, accompanied by error-bars. The best power law fit to the data is provided along with each subfigure, in red. The chromosome number is mentioned in the bottom corner of each subfigure. The range of exponents provided by the fit is 1.09 – 1.40.

**Figure S12:**
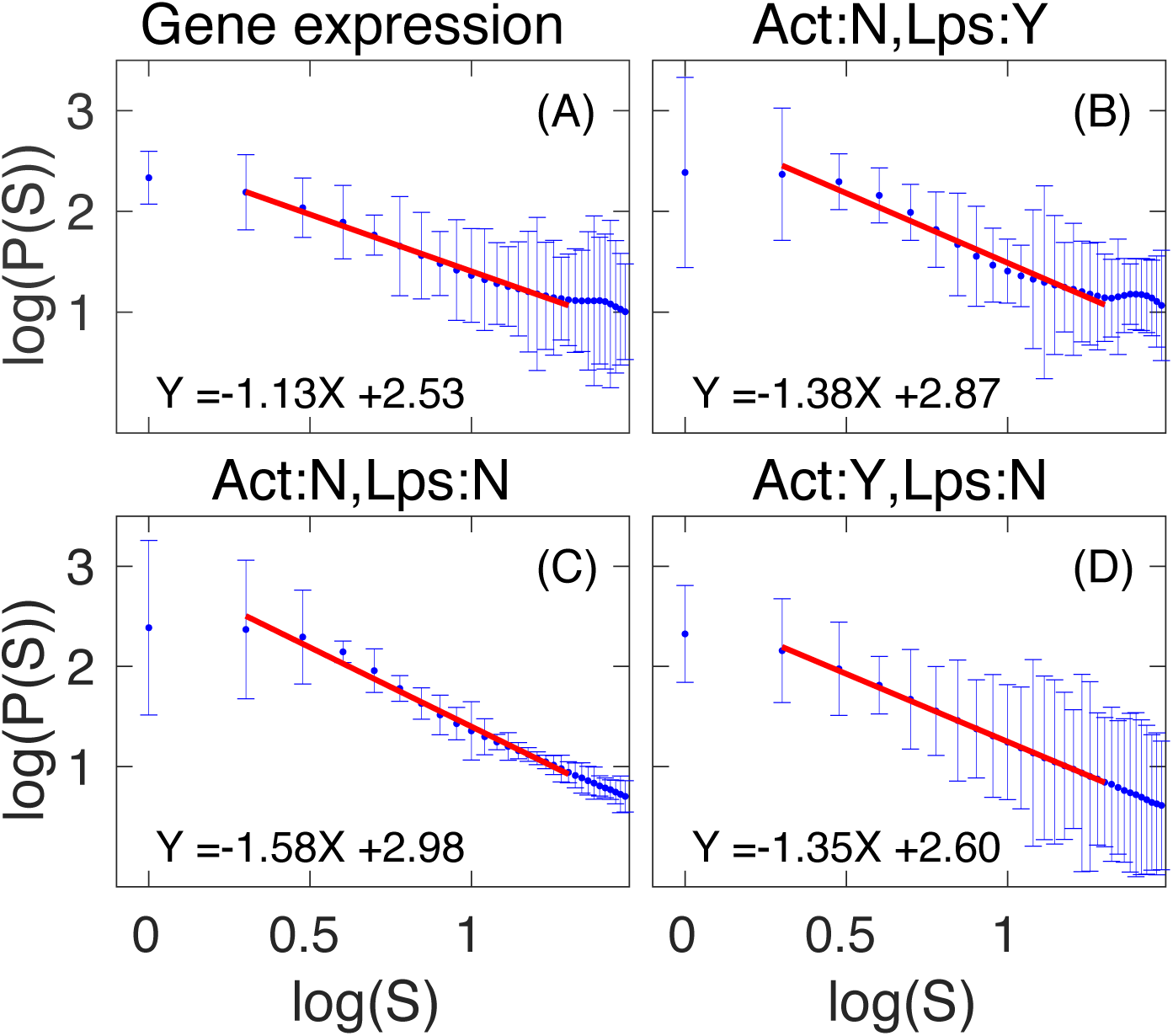
Contact probability P(s) of chromosome 1 for different combinations of activity and loops. Contact probability for chromosome 1, fit to a power law for the gene density, gene expression and combined models, plotted in the range [1-20 Mb]. **(A) Gene Expression** *P*(*s*) for the gene expression model with effective temperature assignments using RNA-seq of GM12878 cell (Consortium et al., 2012). Loops are obtained from Hi-C experiments (Rao et al., 2014); **(B) Act:N, Lps:Y** *P*(*s*) for the case where there is no activity but looping is present; **(C) Act:N, Lps:N** *P*(*s*) for the case where both activity and looping are absent; **(D) Act:Y, Lps:N** *P*(*s*) in the case where the effective temperature assignment corresponds to the combined model for the GM12878 cell type but loops are absent.

**Figure S13:**
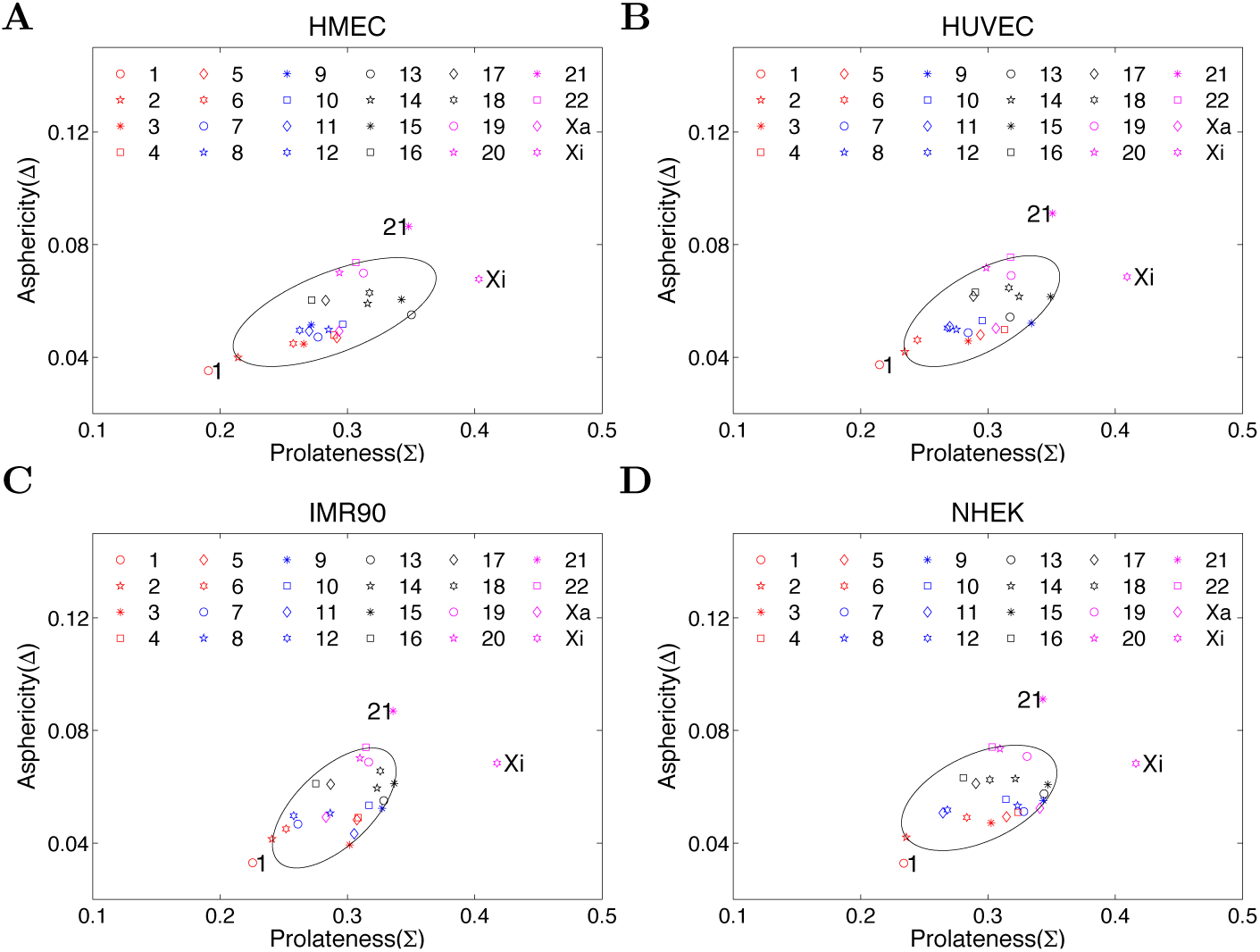
Asphericity Δ and Prolateness Σ parameter of chromosomes for IMR90, HMEC, HUVEC, and NHEK cell type. We capture broad trends in the data by plotting an ellipse that includes closely correlated points. Chromosomes 1, 21 and Xi, are clearly excluded while chromosomes 22, 2, 20, 11, 15, 9 and 13 lie within the ellipse.

**Figure S14:**
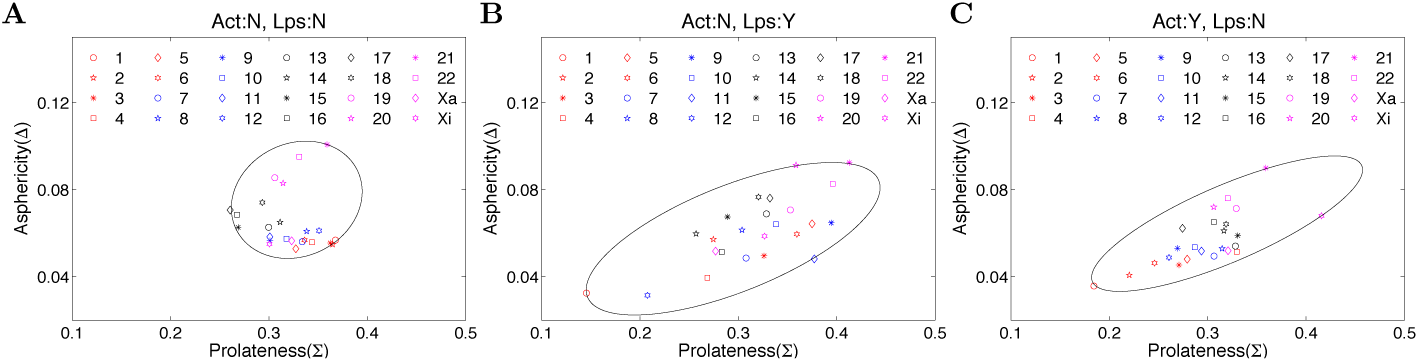
Asphericity Δ and Prolateness Σ parameter of chromosomes for different combinations of activity and loops. Δ and Σ of each chromosome is plotted for 3 different models: **Act:N, Lps:N** in which activity is absent and all permanent loops are absent; **Act:N, Lps:Y** in which activity is absent, but permanent loops are present; **Act:Y, Lps:N** in which effective activity is taken from the combined model appropriate to GM12878 cell type but permanent loops are absent; From these figures, we observe that chromosomes achieve a prolate ellipsoidal shape in the absence of activity. Larger chromosomes are more spherical and smaller chromosomes are rougher and more rod-like.

**Figure S15:**
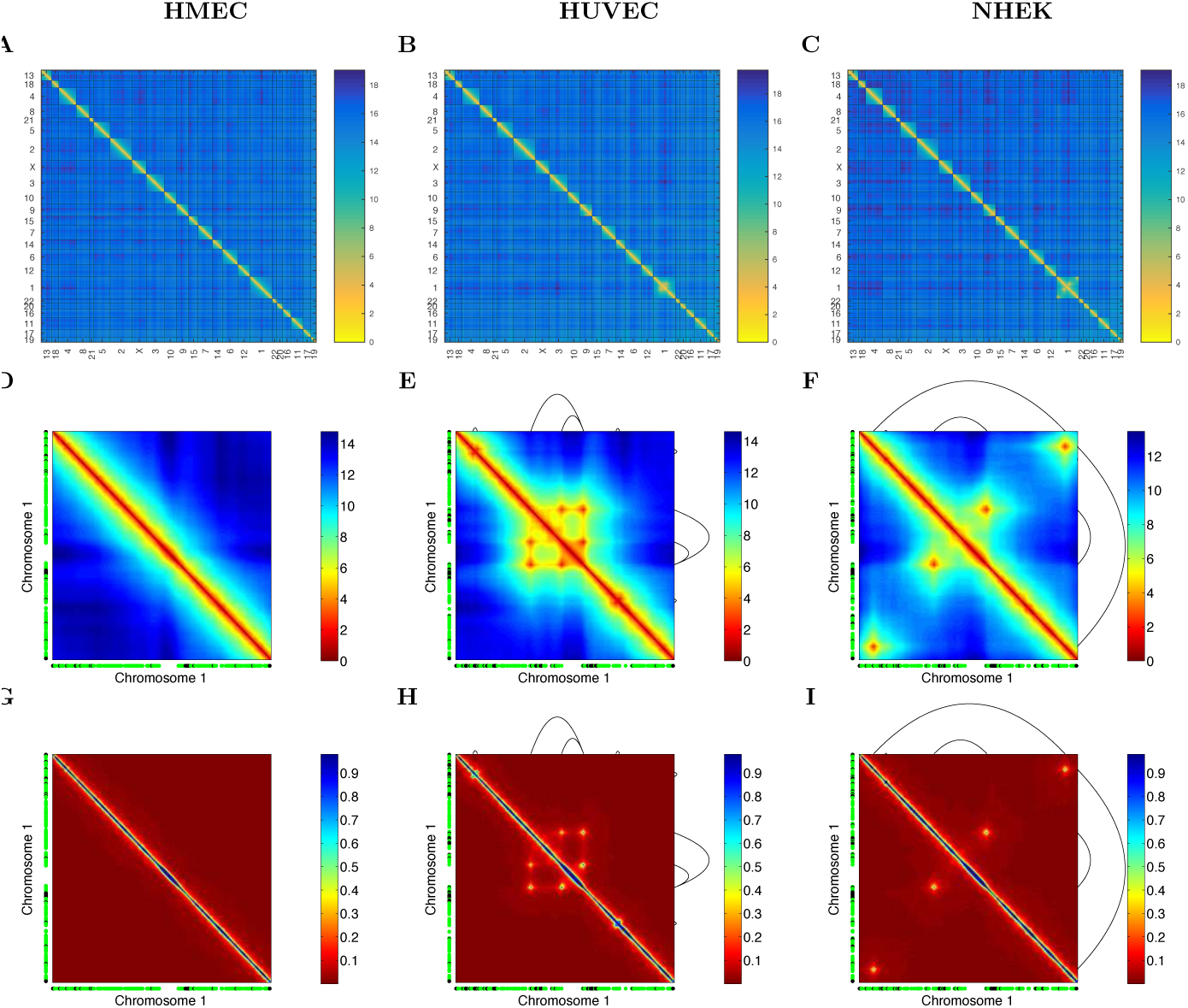
Simulation-derived distance and contact maps for the HMEC, HUVEC and NHEK cell types. **A-C** Distance maps for all chromosomes, ordered according to their gene density; **D-F** Distance map of chromosome 1 displayed along with locations of the permanent loops (black curve) as inferred from the Hi-C data for this cell type. Individual monomers at T = 6 and 7 ≤ *T* ≤ 12 are shown in green and black beside the X and Y axis, respectively; **G-I** Contact map of chromosome 1 displayed along with locations of the permanent loops with black curve as inferred from the Hi-C data. Individual monomers at T=6 and 7 ≤ T≤ 12 are shown in green and black beside the X and Y axis, respectively; Related color-bars for distance and contact maps are shown beside the respective sub-figures.

**Figure S16:**
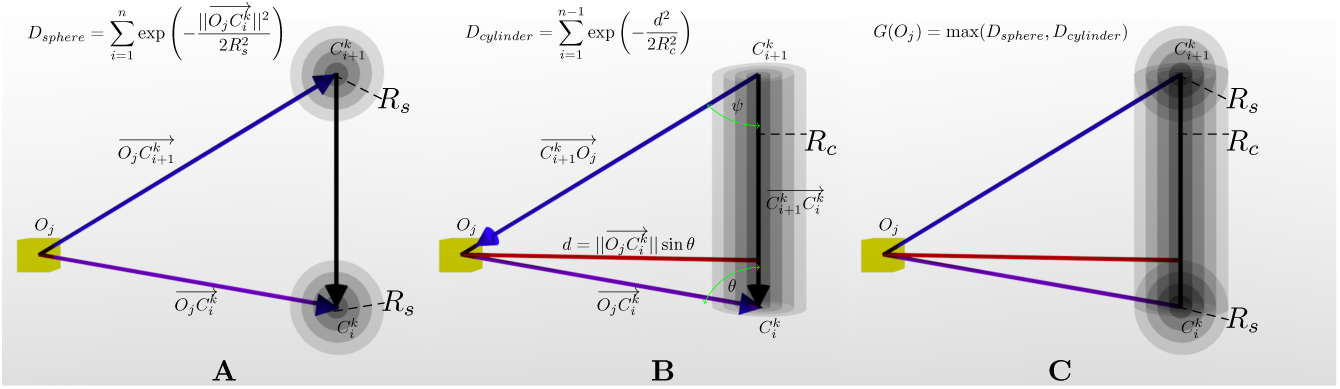
Schematic showing how the chromosome territory is computed using the grid method. **A** Density at yellow grid point (*D_sphere_*) is computed due to sphere effect; **B** Density at yellow grid point (*D_cylinder_*) is computed due to cylinder effect; **C** The actual density at yellow grid point *G*(*O_j_*) is the maximum of either *D_sphere_* or *D_cylinder_*.

**Figure S17:**
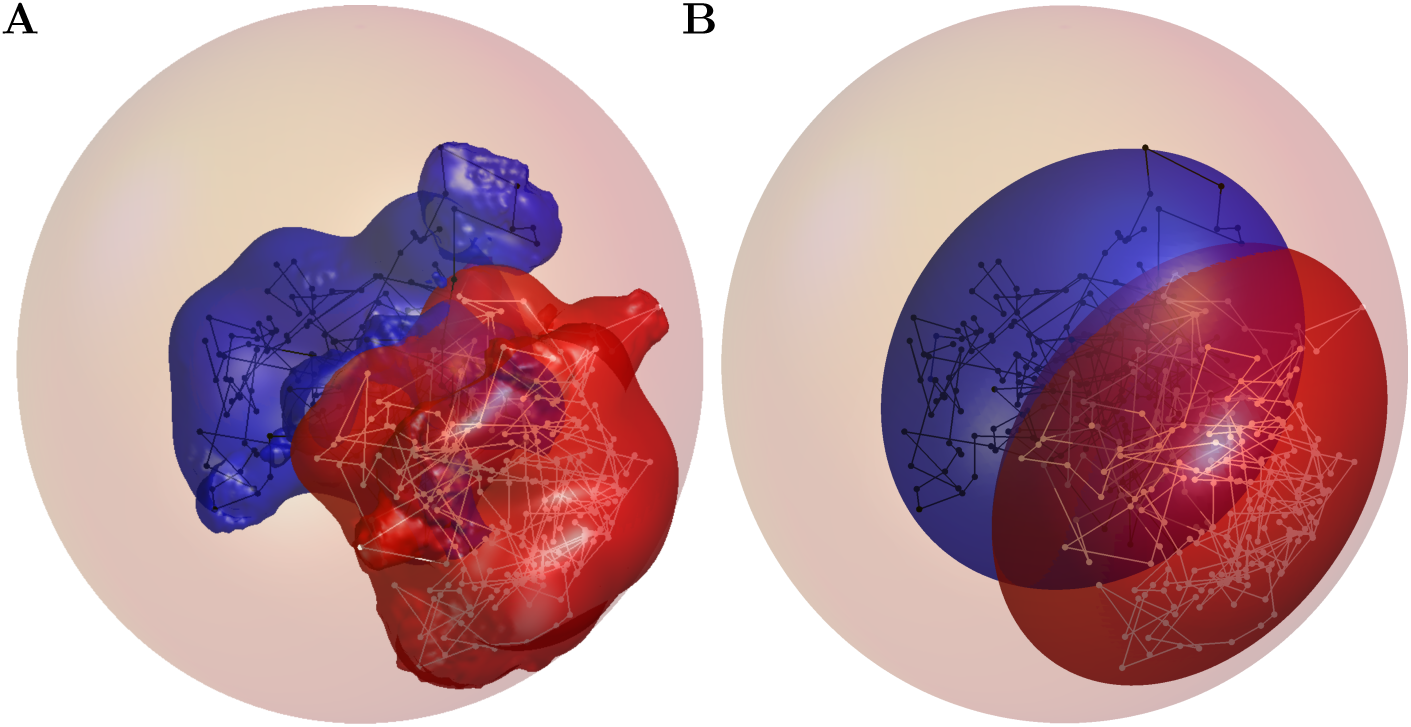
Comparison of output of grid method and 3d ellipsoid fit method. **(A)** Schematic representation of volume overlap of chromosome 1 and 3 by an implicit surface method. The polymer form of chromosome 1 and 3 is represented with white and black color respectively. The 3d surface of chromosome 1 and 3 computed from grid method is shown with red and blue color respectively. **(B)** Schematic representation of volume overlap of chromosome 1 and 3 by an ellipsoidal fit method. The polymer form of chromosome 1 and 3 is represented with white and black color respectively similar to Fig. A. The 3d surface of chromosome 1 and 3 computed from ellipsoid algorithm is shown with red and blue color respectively.

